# Investigating fNIRS Test-Retest Reliability During Lexical Decision

**DOI:** 10.64898/2026.07.14.737938

**Authors:** Laura M. Elliott, Saisha S. Rankaduwa, Cindy Hamon-Hill, Devon A. Bode, Marilla V. M. Hulls, Paige J. A. Lavoie, Aaron J. Newman

**Affiliations:** Department of Psychology & Neuroscience, Dalhousie University

**Keywords:** Functional Near-Infrared Spectroscopy, Test-Retest Reliability, Reading, Lexical Decision

## Abstract

**Significance:** fNIRS is highly suitable for the study of reading development, however, the reliability of its signals is not well understood during reading tasks.

**Aim:** Therefore, this study assessed the test-retest reliability of the fNIRS signal during a common event-related reading paradigm.

**Approach:** English-speaking adults (n = 30) completed a lexical decision task during fNIRS recording twice, one week apart.

**Results:** Our results demonstrated contrast effects partially consistent with prior neuroimaging literature, insofar as for each contrast, at least one predicted region of interest was activated. However, we did not identify significant activation in all predicted brain areas. Regarding group level test-retest reliability, we observed poor reliability across predicted brain regions for most conditions, with the exception of fair test-retest reliability in the left posterior temporal lobe for coarse lexical tuning. At the single-subject level, test-retest reliability ranged from poor to excellent across subjects, but was poor for most subjects.

**Conclusion:** These results suggest fNIRS can detect changes in brain activation during a fast event-related reading task at the group level. However, reliability does not appear sufficient to interpret individual-level data. Further research should explore reliability across a wider range of designs to assess the generalizability of these findings.

## Introduction

Functional near infrared spectroscopy (fNIRS) is a non-invasive, functional neuroimaging technique with adequate spatial and temporal resolution for cognitive neuroscience research.^1–3^ fNIRS has many benefits relative to other non-invasive neuroimaging techniques, such as being relatively insensitive to head motion, silent, and comfortable for the participant. ^2,3^ However, the reliability of the fNIRS signal requires further research. It is important to establish the reliability of fNIRS measurements at the group level to ensure that results are likely to be reproducible. Further, if fNIRS is to be used to investigate changes in brain activity longitudinally, or to relate individual differences in brain activity to behavioral abilities, it is necessary to confirm fNIRS provides a reliable snapshot of task-related brain activity at the level of individual participants. While fNIRS has been deemed reliable at the group level in select tasks (reviewed below), its reliability has not been assessed with reading tasks. At the single subject level, reliability is also not yet well established and has often been reported as being low.^1,4^

Therefore, the first objective of the present study is to establish the test-retest reliability of the signals recorded by fNIRS during an event-related reading paradigm (lexical decision task), at the group and single subject level. Additionally, as sufficient work has not yet been done to validate fNIRS in the study of reading,^5,6^ the second objective is to identify the brain areas that demonstrate significant activation for the effects of interest during the reading task, as measured by fNIRS, and identify whether these are consistent with results of prior research using fMRI.

## Reliability of the fNIRS Signal

Test-retest reliability is the consistency between two measurements of the same thing, taken at different time points,^7^ and can be assessed at the group and single subject levels. A significant proportion of existing neuroimaging research relies on group-level reliability, which can be adequate in the absence of subject level reliability.^8,9^ However, to investigate changes in brain activity over time, the neuroimaging technique also needs adequate subject level reliability, to ensure changes observed over time are greater than the error introduced by the measurement technique.^7,8,10^ Further, to investigate individual differences in brain activity and relate those differences to variables of interest (e.g., behavioral abilities), adequate subject level reliability is required.

A common measure for assessing test-retest reliability is the intraclass correlation coefficient (ICC), which directly assesses agreement across multiple measurements of the same thing using analysis of variance to calculate the error variance due to the measurement tool.^7,8,10–13^ ICC provides a score ranging from 0-1, with common criteria for interpretation used first in psychology research, and then later in neuroimaging studies, being excellent (> 0.75), good (0.59 to 0.75), fair (0.40 to 0.58), and poor (< 0.40).^8,14^

The test-retest reliability of the signals recorded using fNIRS has been investigated across a variety of tasks, with test-retest reliability ranging from fair to excellent at the group level across studies, task designs, and brain regions.^4,9,15–18^ Even within a given domain, such as motor tasks, test-retest reliability at the group level has been demonstrated to differ by task type within the same group of participants,^19^ as well as between replications of the same task.^15,17^ While limited research has assessed test-retest reliability of the fNIRS signal during language processing, the reliability of the fNIRS signal has been shown to be fair at the group level during both a verbal fluency task,^18^ and passive listening.^9^ Greater reliability is observed when averaging data across a number of fNIRS channels covering a region of interest (ROI), as compared to when analyzing data at the single channel level.^4,9,18^

Only a small subset of studies have investigated the reliability of the fNIRS signal at the subject level.^4,16,18,20^ While some participants demonstrate good to excellent test-retest reliability of task-related signals, others demonstrate minimal task-related activity, or task-related activity that differs from one session to the next, resulting in poor reliability.^18,20^ A number of factors have been identified, or hypothesized, to contribute to poor intra-individual reliability across fNIRS sessions.

One suggested factor is participant state, such as affect or drowsiness, differing from one study visit to the next.^9,16^ Sleep deprivation and self-reported sleepiness are also associated with changes in the location and magnitude of the hemodynamic response in the brain.^21,22^ On the other hand, Huang et al. (2017) assessed participants’ affective states prior to each session and found no significant differences from one session to the next in participant affect, yet still observed poor subject-level reliability across sessions. Additionally, moderate physical activity prior to neuroimaging results in both increases and decreases in cerebral blood perfusion, with the change in cerebral blood flow varying by brain region.^23^ Variability in the fNIRS signal at the subject level has also been suggested to vary with task type. For example, less-structured tasks may lead to more variable performance across repetitions of the task, and thus more variable task related brain activity from one session to the next.^18^ Hair colour and thickness may also contribute to differences in subject-level reliability between individuals, as both have been demonstrated to influence fNIRS signal quality.^24,25^ Additionally, it has been speculated that session-to-session variability in fNIRS optode placement on the scalp may reduce the reliability of the signal, though this variability has not been quantified.^19^

Pharmacological substances with vascular action directly influence cerebral blood flow and consequently the signal measured by fNIRS.^26^ Of the pharmacological agents, caffeine is among the most prevalent, and has been shown to influence both neural activity and blood flow in the brain.^27^ The influence of caffeine on physiology is complex, as its modulation of cerebral blood flow varies across brain regions, and is dependent on the individual’s habitual caffeine intake.^28,29^

Overall, it is clearly necessary to consider variables such as caffeine consumption, sleep, and physical activity when investigating the test-retest reliability of fNIRS. As well, since task can affect reliability, it is important not to assume that reliability metrics from one experimental paradigm will necessarily generalize to another setting.

## The Lexical Decision Taks and fNIRS

The lexical decision task is a widely used experimental paradigm for studying the processes involved in single word reading.^30^ In a lexical decision task, participants read word or word-like stimuli and indicate if they are reading a real or non-real word. Performance during lexical decision relies on the engagement of orthographic, phonological, and semantic information ^31^. While it has been used extensively in electroencephalography (EEG) and fMRI studies, it has received limited investigation with fNIRS (e.g., Hofmann et al., 2008; Sela et al., 2014; Stela et al., 2011), and no studies have examined its reliability in this context.

Among the commonly obtained contrasts from lexical decision tasks, print tuning is an increased neural response to letter or text stimuli as compared to that for meaningless symbols; isolating orthographic processing. Using fMRI, print tuning has been associated with activation in the left parietal cortex, superior and inferior frontal gyri, fusiform gyrus, and inferior occipital-temporal regions.^34–36^

Sublexical tuning is the contrast between pronounceable non-words (“pseudowords”, e.g., “blorp”) and unpronounceable patterns of letters (i.e., consonant strings), isolating processes involved in converting orthographic information to phonologic representations. fMRI studies of word reading have associated the left anterior superior temporal gyrus,^37^ supramarginal gyrus,^38^ inferior frontal gyrus,^39–41^ angular gyrus,^41^ and inferior parietal lobule^39^ with sublexical tuning.

Lexical tuning is the contrast between real words and non-words. Lexical tuning contrasts can be described as “coarse” or “fine”, depending on whether the words are pronounceable or not. Coarse lexical tuning is the contrast between real words and non-pronounceable strings of letters (i.e., consonant strings) and fine lexical tuning is the contrast between real words and pseudowords. In fMRI studies, coarse lexical tuning is associated with increased activation in the visual word form area, left posterior temporoparietal cortex and left inferior temporal cortex,^42,43^ while fine lexical tuning is associated with activation in the left angular, anterior superior temporal, and middle temporal gyri, and the left inferior and superior parietal lobule.^44–49^

Across previous fNIRS studies using the lexical decision task, only fine lexical tuning has been investigated — in German,^33^ Hebrew,^30,32^ and Mandarin^50^ — and results are inconsistent both between studies, and with prior fMRI work. Words have been reported to elicit greater activation than pseudowords in the left superior frontal and inferior parietal gyri,^33^ and the left upper- and mid-frontal lobe,^30,32^ while the third study reported activation only for pseudowords more than real words, in the left inferior frontal gyrus.^50^ The paucity of evidence from fNIRS, and the inconsistencies both between fNIRS studies and with prior fMRI results, underscores the need for further validation of the lexical decision task using fNIRS, and investigation of the reliability of the results at the individual and group levels.

## The Present Study

In the present study, we assessed the test-retest reliability of the neural activity for print tuning, sublexical tuning, coarse and fine lexical tuning contrasts as measured by fNIRS, at both the subject and group levels. We recruited adults to complete two identical study visits, one week apart, and had participants report on variables that may affect the fNIRS signal at the outset of each session.

The first objective of this study was to assess the test-retest reliability of the fNIRS signal during lexical decision. Based on prior literature, we hypothesized the intraclass correlation (ICC) would show good (0.59 < ICC < 0.75) reliability at the group level,^4,9,16,18^ with greater reliability when data was analyzed across clusters of channels (i.e., region of interest level) as compared to when data was analyzed at the individual channel level.^4,9,18^ Additionally, at the individual participant level, we hypothesized we would observe fair (0.40 < ICC < 0.59) reliability for the majority of participants.^9,18^

The second objective was to identify which brain regions demonstrate significant activation for the effects of interest as measured by fNIRS, and if these results align with prior fMRI literature. We hypothesized we would observe a significant print tuning effect (i.e., increased brain activity for consonant string as compared to false font stimuli) in the left inferior parietal lobe (LIPL) and inferior frontal gyrus (LIFG).^34–36,51^ While a significant print tuning effect has also been observed in the left fusiform gyrus and left inferior occipito-temporal regions,^34^ these brain areas cannot be reached by fNIRS due to the limited penetration depth of the near-infrared light.^2^ We also hypothesized a significant sublexical tuning effect (i.e., increased brain activity for pseudoword as compared to consonant string stimuli) would be observed in the left anterior temporal lobe (LATL),^37^ LIFG, ^39–41^ and LIPL.^39^ We hypothesized a significant coarse lexical tuning effect (i.e., increased brain activity for real words as compared to consonant string stimuli) would be observed in the left posterior temporal lobe (LPTL) and LIPL.^42,43^ Finally, we hypothesized a significant fine lexical tuning effect (i.e., increased brain activity for real words as compared to pseudoword stimuli) would be observed in the LIPL and the LATL.^44–46^

## Methods

### Participants

The minimum sample size for a reliability study with two measurement instances, with ICC[H_0_] = 0.39, ICC[H_1_] = 0.75, α = 0.05, power = 0.80, was computed to be 21 participants.^52,53^ Thirty-three English-speaking adults were recruited through the participation system for Dalhousie University students. Participants were recruited beyond the required sample size to account for potential attrition and data loss due to poor signal quality or technical issues. Of the 33 participants, data from two participants was excluded due to poor signal quality (defined below), and data from one participant was excluded as they only completed one session. The final sample of 30 participants consisted of 13 males and 17 females, ranging in age from 18 to 37 years (mean age = 21.47, SD = 4.8). Based on the Edinburgh Handedness Inventory – Short Form,^54^ 28 participants were right-handed, one was left-handed, and one had mixed handedness. Participants were not excluded due to handedness.^55^

While all participants could read and speak English fluently, five participants learned English as their second language. In total, 18 participants spoke two languages, and 6 of those participants spoke a third language. All participants had normal or corrected-to-normal vision, no self-reported neurological conditions (e.g., epilepsy), and did not use medications affecting brain activity (e.g., anti-seizure medications). Participants provided signed informed consent, according to the Declaration of Helsinki, at the beginning of the first study visit, and were reminded their participation was voluntary at the subsequent visit. Participants received points redeemable for course credit as compensation. This research was approved by the Dalhousie University Social Sciences and Humanities Research Ethics Board (#2023-6928).

## Materials

### Questionnaire

We collected demographic information via electronic questionnaire including age, gender, and language history, and assessed handedness using the *Edinburgh Handedness Inventory – Short Form.*^54^ This questionnaire included the *PANAS-X Positive and Negative Affect Schedule – Expanded Form*, which has been deemed an appropriate measure of current affect.^56^ Additional questions asked participants about their sleep the night prior, recent caffeine consumption and physical activity. This questionnaire was used in Session 1, and again in Session 2 without repeating demographic questions.

### Lexical Decision Task Stimuli

Four types of word stimuli were used in the lexical decision task: real words, pseudowords, consonant strings, and false fonts. The 60 real words were selected from a list of nouns that were six characters in length or less, from reading outcome word lists for the first, second, and third grade, to ensure they were readily recognizable. The 60 pseudowords were generated by a natural language model that combined high frequency pairs of letters from the English language, to form items matched in length to the list of real words, that follow legal letter patterns. The 60 consonant strings were developed by replacing the vowels in each real word with randomly selected consonants. Finally, the 60 false font stimuli were created by presenting the 60 pseudowords in the *Brussels Artificial Character Sets* (BACS) font.^57^

The study included an active baseline task during intervals between word stimuli. The set of 36 unique active baseline screens consisted of eight cartoon animals on one of six nature backgrounds. On each screen, there were two animals at the top, two to the right and two to the left of center, and two at the bottom of the screen.

## Procedure

Participants completed two sessions following the same procedure, on the same day of the week and at the same time of day, one week apart. Each session took place at Dalhousie University, and took approximately one hour. Sessions were conducted by the same pair of researchers for each participant, and the researchers followed a script when interacting with the participant, to ensure consistency across sessions.

### Sessions 1 & 2

Upon arrival at the lab, signed informed consent (Session 1) or continued verbal assent (Session 2) was obtained.

Participants were fitted with a stretchable fabric cap (EasyCap, n.d.) prepared with 16 light emitters and 16 detectors (optodes), forming a 42-channel system, in positions according to the international 10-20 system.^58^ The interoptode distances for all emitter-detector pairs were 3 to 4 cm. The montage had eight short-separation channels (located at FC6, T8, CP6, P8, FC5, T7, CP5, and P7) to measure non-neural physiological noise, and one accelerometer on top of the head to monitor head movements. The montage, shown in Figure 1, was optimized using NIRSite (v2.0, NIRx) to cover the inferior frontal, superior and middle temporal, and inferior parietal cortices of both cerebral hemispheres. Data was collected using a NIRSport2 fNIRS system (NIRx, n.d.), which employs continuous-wave measurement using LED emitters with dual-tip optodes that shine near-infrared light at two wavelengths (760 nm & 850 nm), with a sampling rate of 6.104, and a maximum light intensity of 25mW per wavelength of light.

**Figure 1.**
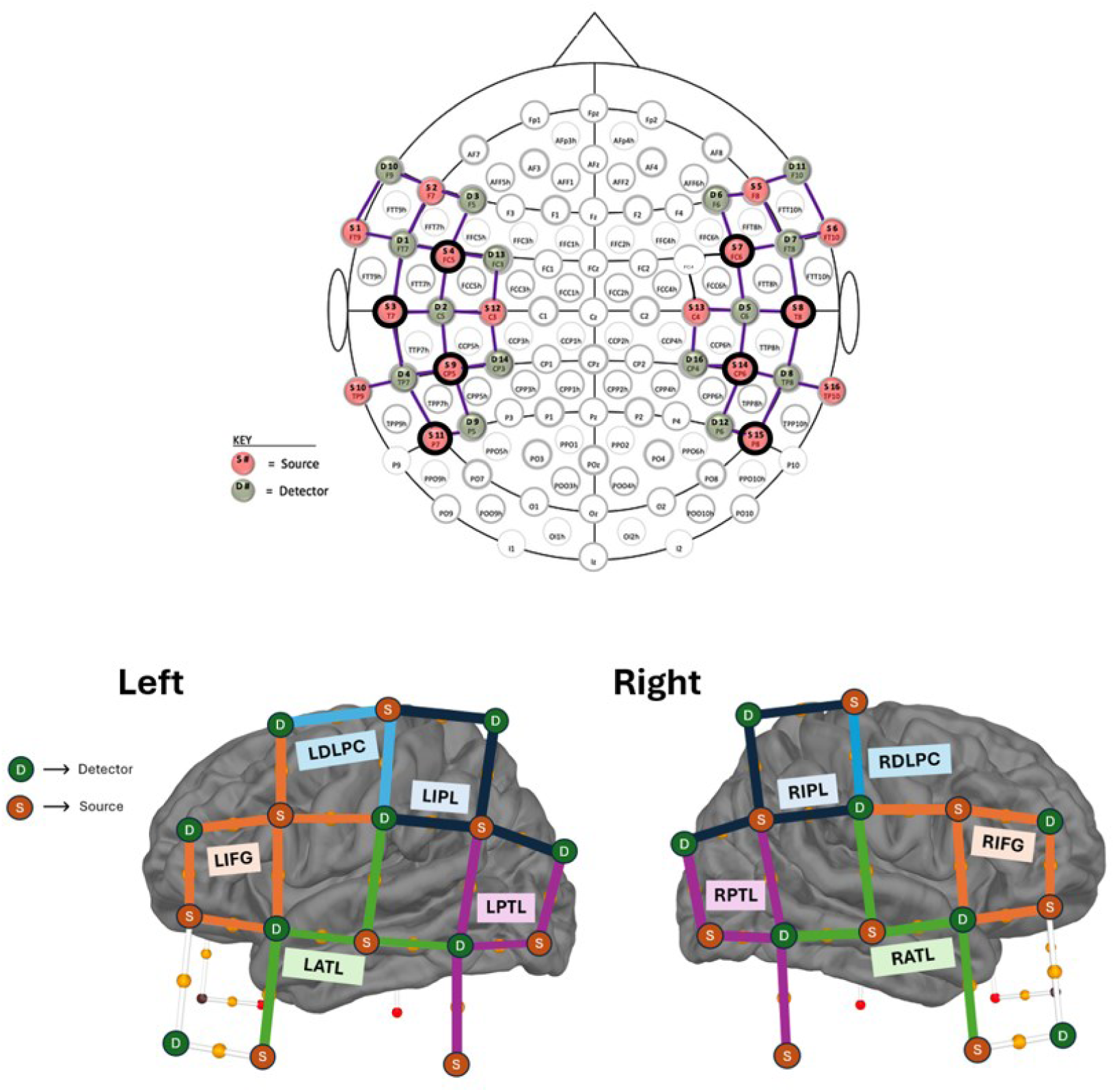
fNIRS Montage Illustration. *Note.* Illustration of the fNIRS montage of 16 light emitters and 16 light detectors, (Upper) according to the international 10-20 system (Acharya & Acharya, 2019), with black outlined sources indicating the location of short-separation channels, and (Lower) the channels contributing to each region of interest investigated in the left hemisphere and right hemisphere. Both images were adapted from NIRx NIRSite 2.0, https://nirx.net/nirsite.

Once the fNIRS cap was in place, the participant completed the questionnaire. Then, participants placed their chin in a chin rest 47.5 cm away from a Dell monitor (27-inch, 1920 x 1080 resolution, VG2722HS). Next, light emitter optimization signal quality checks were performed using Aurora (v2021.4, NIRx), and necessary adjustments to optodes were performed. The lighting in the window-less room was dimmed to 50%, to reduce ambient light.

Then, fNIRS recording began and participants completed a practice run of the lexical decision task. The task was presented using the presentation software PsychoPy (v2022.2.5).^59^ Participants were provided a standard video game controller, and the initial screen of the task provided participants instructions for which buttons to press.

During the practice, participants were presented each word type twice in pseudorandomized order, using stimuli matched to those in the main task. Then, participants completed five runs of the lexical decision task. During each run (6 minutes and 18 seconds in length), participants were randomly presented 36 real words (e.g., STAR), 12 pseudowords (e.g., SPET), 12 consonant strings (e.g., STLR), 12 false fonts, and 36 active baseline screens. The presentation of the stimulus words was organized such that after all five runs, participants had been shown each real word three times, creating 180 real words, 60 pseudowords, 60 consonant strings, 60 false fonts, and 180 active baseline screens.

Across both the practice and lexical decision task, real words, pseudowords, and consonant strings, were presented in the font “KG Neatly Printed” with a letter height of 1.5 cm (visual angle of 1.809), and all words were presented in black on a medium gray background. Each word and baseline search task was on screen for 2000 ms. A fixation cross was presented for 1500 ms after each stimulus. The order of the presentation of the stimuli, and the number of sequential repetitions of each (e.g., three active baseline tasks in a row, then two real words in a row), was determined by an algorithm to maximize fNIRS signal contrast between conditions.^60^

## Data Analysis

### Behavioral Data

The negative affect, positive affect, fatigue, and attentiveness index scores on the PANAS-X^56^ were calculated and analyzed in Python (v3.12.13), for each participant and study visit. Participants rated their caffeine consumption and sleep quality as “below average”, “average”, or “above average” based on what is typical for them, and sleep quantity was reported as “less than 6 hours”, “between 6 and 8 hours”, “more than 8 hours”. The Wilcoxon Signed-Rank test was performed on each variable, to identify significant differences between sessions 1 and 2, using the SciPy package.^61^

Accuracy and reaction time for the lexical decision task were analyzed using the lme4 package^62^ in R (v4.4.2). For accuracy, single-trial data (correct/incorrect) was analyzed using linear mixed effects with a binomial family. We tested four models with condition as a fixed effect, with and without session as a fixed effect, and with random effects for participants, stimulus item, and stimulus condition (i.e., word type). The best model was selected on the basis of Akaike’s information criteria (AIC) weights,^63^ and included fixed effects for condition and session, with random effects for participants and stimulus item. For reaction time, we also used linear mixed effects modelling, but with a Gaussian family to account for the non-normal distribution of reaction times.^64^ We tested four models, all with fixed effects for condition and session, with random effects for participants, stimulus item, and stimulus condition. Similarly, the best model was selected using AIC weights,^63^ and included fixed effects for condition and session, with random effects for participants.

### fNIRS Data

#### Preprocessing

fNIRS data was preprocessed using the MNE (v1.9.0)^65^ and MNE-NIRS (v0.7.1)^65,66^ software packages in Python (v3.12.8). First, raw intensity values were converted to optical density. Then, optical density data was assessed for channels with poor scalp coupling, and channels with a scalp coupling index below 0.7 were marked as ‘bad’ channels. Next, temporal derivative distribution repair was used to identify and remove motion artifacts, (baseline shift and spike artifacts). Short channel regression was then performed to reduce the influence of physiological signals on the data. Next, optical density data was converted to hemoglobin concentration using the modified Beer-Lambert law, and a negative correlation enhancement algorithm was used to further reduce the influence of motion artifacts. The data was then low pass filtered with a cutoff of 0.5 to remove physiological (cardiac and respiratory) artifacts. Lastly, channels marked as ‘bad’ were interpolated, such that data at the marked channels was recreated by interpolating the signal from nearby acceptable channels.

#### First Level Analysis

For each experimental run, a design matrix was created to model the expected neural response to the presentation of each stimulus, convolved with a gamma function.^67^ Additional regressors were included in each design matrix to account for known sources of variance, including the temporal derivative of each condition’s gamma functions; the 8 fNIRS short channels; the six largest principal components derived from the nine raw accelerometer values, and a series of cosine basis functions up to 0.01 Hz to model low frequency artifacts. This design matrix was used to fit a general linear model to the preprocessed data using the MNE-NIRS library.

#### Second Level (Group) Analysis

The parameter estimates from the first-level analyses for each run were imported into R (v. 4.4.2) and two data cleaning steps were performed. First, we computed the standard deviation of the number of channels rejected across all runs for all participants, and a threshold of >2 standard deviations was used to reject runs with an excessive number of bad channels. Second, outlier values were removed by converting the parameter estimates within each participant, session, and chromophore to *z* scores and then removing any values +/- 2 standard deviations from the mean. Linear mixed effects modelling, implemented in the mgcv library (v. 1.9.1),^68^ was then used to analyze the first-level coefficient (theta) values of each chromophore (HbO or HbR), with session, condition, ROI, and chromophore as fixed effects, and random intercepts for channel by subject, for run by session, and for session by subject. The estimated marginal mean (computed using the emmeans package, v.1.10.7) for each chromophore was plotted by word type and brain region. Then, pairwise contrasts of the estimated effect size for each chromophore were calculated in each ROI, for each contrast of interest (i.e., print tuning, sublexical tuning, and lexical tuning), both across sessions, and for sessions 1 and 2 separately. The resulting *p* values were corrected for the number of contrasts using the false discovery rate method, and significance was determined using α = 0.05.

## Test-Retest Reliability Analysis

### fNIRS Data

To investigate test-retest reliability of the fNIRS signal, estimates of each chromophore (HbO or HbR) also needed to be calculated at the subject level. Therefore, linear mixed effects was used to analyze the coefficients of each chromophore (HbO or HbR) at the subject level, with condition, channel, session, and chromophore as fixed effects, and random intercepts for channel within session. Then, pairwise comparisons were computed between word types at the subject level, to calculate the magnitude of activation for each contrast, at each channel, for each session. The resulting *p* values were corrected for the number of contrasts using the false discovery rate method.

Data was then aggregated at the subject level at both the channel and the ROI level (clusters of channels). To obtain channel level test-retest reliability data, the estimated marginal means at each channel, by session, were used as inputs. For the ROIs, estimated marginal means were averaged across clusters of channels corresponding to pre-defined ROIs, for each session.

Additionally, data was aggregated at the group level at both the channel and ROI level. To obtain group level test-retest reliability data, the same procedure for averaging estimated marginal means was used, with the addition of averaging the results across subjects, for session one and two. All intraclass correlations were calculated using the averaged estimated marginal means and the Pingouin package (v0.5.4) in Python (v3.12.8), to obtain the ‘ICC2’ value, which is a measure of absolute agreement between repeated measures.^69^

### Subject Level Test-Retest Reliability and Behavioral Measures (Exploratory)

To explore the relationship between behavioral data and subject level test-retest reliability, we calculated the absolute difference between scores on relevant behavioral variables measured through questionnaires at the beginning of each session. These variables included self-reported sleep quality, sleep quantity, caffeine consumption, as well as scores on the negative affect, positive affect, attentiveness, and fatigue indices on the PANAS-X.^56^ For each behavioral variable, we then fit a linear mixed effects model using the mgcv package (v1.9.3) in Python (v3.12.8), with subject level ICC as the dependent variable, fixed effects of region of interest, chroma, contrast, and the difference score of the behavioral variable, as well as the four-way interaction of these variables, and random intercepts for subjects.

## Results

### Behavioral Data

#### Questionnaire Data

On the PANAS-X^56^ we observed a significant decrease in positive affect (*W* = 54.0, *p* = .001) and attentiveness (*W* = 45.5, *p <* .001) at session 2 as compared to session 1. No significant difference was observed in negative affect (*W* = 92.5, *p* = .268) or fatigue (*W =* 111.0, *p* = .263), between sessions. Mean PANAS-X index scores are visualized in Supplementary Results, Figure S1. With respect to sleep, there was no significant difference in the amount (*W* = 10.5, *p* = .527), nor in the quality (*W* = 51.0, *p* = .317) of sleep reported prior to sessions 1 and 2. There was no significant difference observed in caffeine consumption between the two sessions (*W* = 16.0, *p* = .763). Distributions of self-reported sleep quantity, quality, and caffeine consumption are visualized in Supplementary Results, Figure S2.

### Lexical Decision Task

Behavioral data from the lexical decision task was trimmed prior to analysis to remove outliers. We removed trials where reaction time was shorter than 150 ms, as this was considered too fast to represent a real decision. Then, reaction times were converted to *z* scores for each participant, and trials with reaction times that were +/- 2.5 standard deviations from the individual’s mean reaction time were removed. This removed 617 trials, or 2.93% of the original data. The trimmed data was used to analyze accuracy and reaction time.

#### Accuracy

Group level mean accuracy for each word type, by session, are visualized in Supplementary Results, Figure S3. Participants demonstrated high accuracy for consonant string stimuli in session 1 (*M* = .998, *SD* = .0420) and 2 (*M* = .999, *SD* = .0238), real word stimuli in session 1 (*M* = .986, *SD* = .116) and 2 (*M* = .996, *SD* = .0630), and false font stimuli in session 1 (*M* = .982, *SD* = .133) and 2 (*M* = .980, *SD* = .140). Accuracy was slightly lower for pseudoword stimuli in both sessions 1 (*M* = .966, *SD* = .181) and 2 (*M* = .969, *SD* = .0630).

Planned pairwise contrasts between sessions 1 and 2 showed significantly greater accuracy for real words in session 2 as compared to session 1 (Estimate = -1.28, *z* = -4.96, *p* < .001). No significant difference in accuracy for false font, consonant string, or pseudoword stimuli was observed between sessions 1 and 2.

#### Reaction Time

Group level mean reaction times for each word type, by session, are visualized in Supplementary Results, Figure S4. In both session 1 and 2, reaction time was fastest for consonant strings (Session 1, *M* = .464 s, *SD* = .256; Session 2, *M* = .417 s, *SD* = .168), followed by false fonts (Session 1, *M* = .492 s, *SD* = .288; Session 2, *M* = .442 s, *SD* = .209), then real words (Session 1, *M* = .497 s, *SD* = .268; Session 2, *M* = .464 s, *SD* = .195), and slowest for pseudowords (Session 1, *M* = .622 s, *SD* = .285; Session 2, *M* = .550 s, *SD* = .230).

Pairwise contrasts showed significantly faster reaction times in session 2 as compared to session 1 for all conditions: consonant string (Estimate = -.580, *z* = -4.54, *p* < .001), false font (Estimate = -.599, *z* = -5.18, *p* < .001), real word (Estimate = -.228, *z* = -3.62, *p* < .001), and pseudoword (Estimate = -.452, *z* = -5.70, *p* < .001) stimuli.

### fNIRS Data

Preprocessed fNIRS data was analyzed at the group level to identify task-related activation. For each contrast, we first analyzed changes in HbO and HbR across both sessions, visualized in Figure 2, to achieve the most sensitivity for detecting a significant effect. Next, we analyzed changes in HbO and HbR for session 1 and 2 separately, visualized in Figure 3. Effect size, confidence intervals, and significance levels for each contrast of interest, both across sessions and for session 1 and 2 separately, are presented in Table 1.

**Figure 2.**
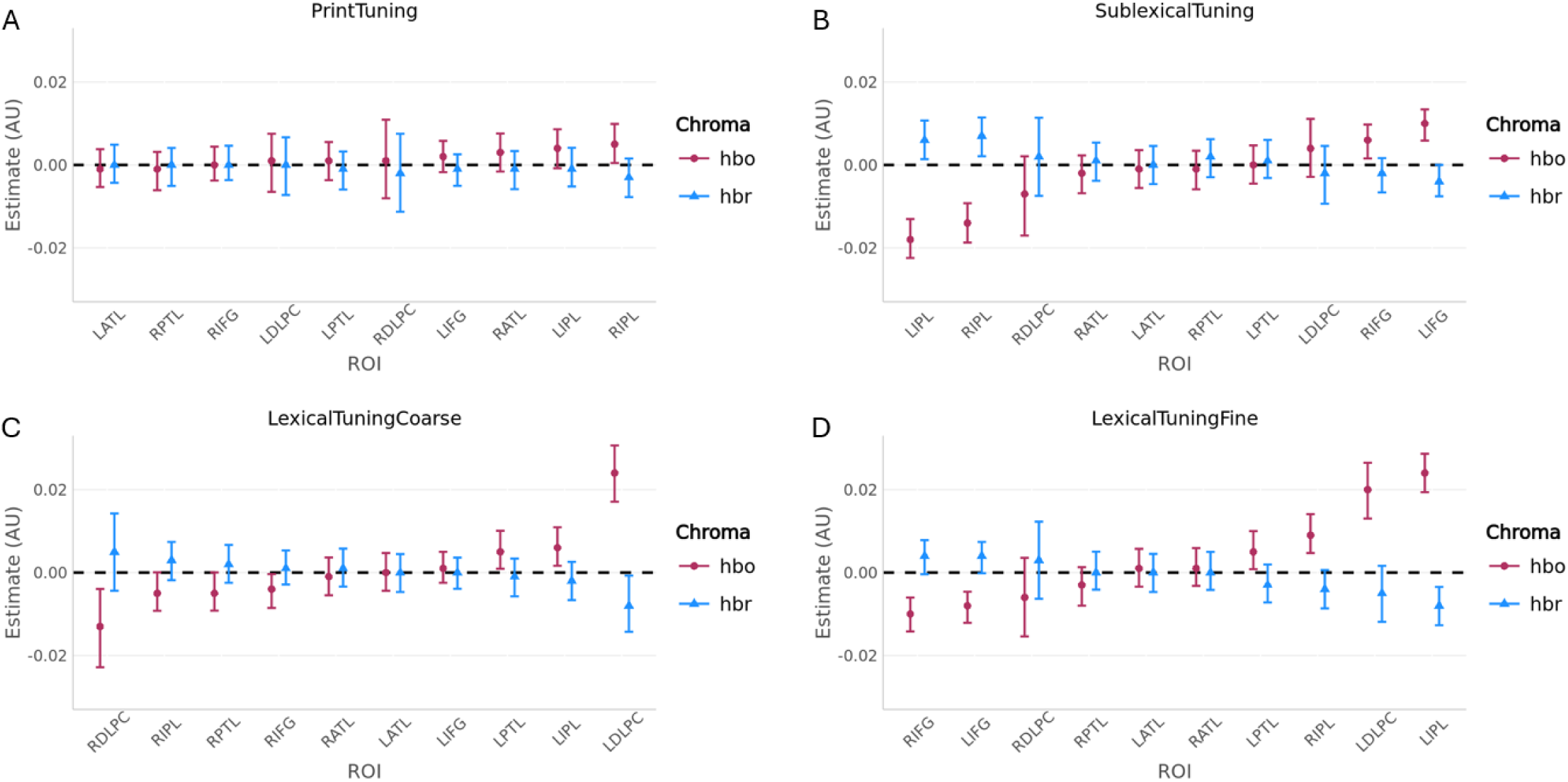
Group Level Estimated Marginal Means for Contrasts of Interest. *Note.* The estimated concentration change of oxygenated (HbO) and deoxygenated hemoglobin (HbR) at the group level, when data is averaged across sessions, for (A) the print tuning contrast (consonant string stimuli minus false font stimuli), (B) the sublexical tuning contrast (pseudoword stimuli minus consonant string stimuli), (C) the coarse lexical tuning contrast (real word stimuli minus consonant string stimuli), and (D) the fine lexical tuning contrast (real word stimuli minus pseudoword stimuli). In each panel, the regions of interest (ROI) are sorted such that the ROI showing the largest HbO effect is on the right and the smallest HbO effect is on the left.

**Figure 3.**
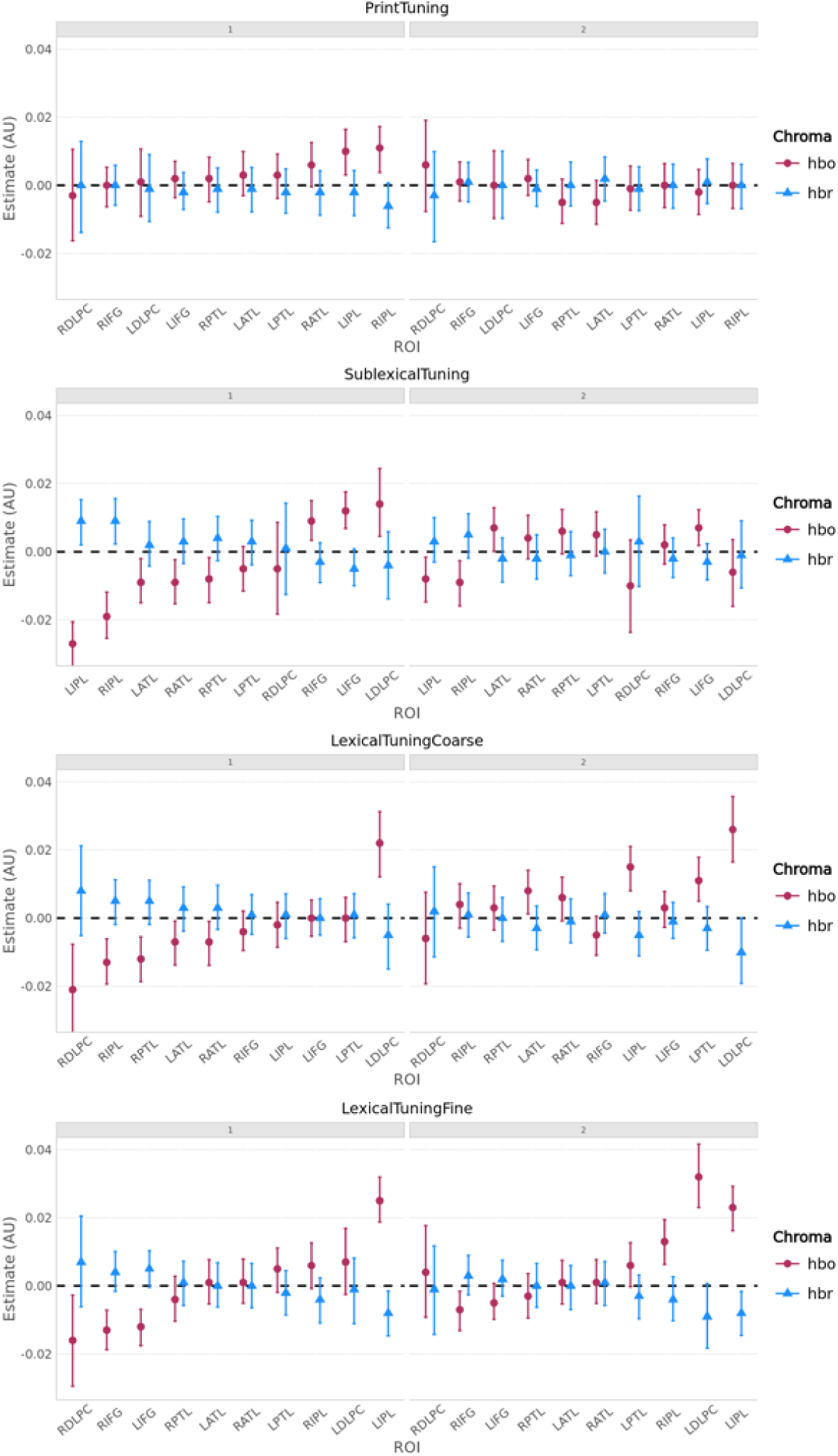
Group Level Estimated Marginal Means for Contrasts of Interest, by Session. *Note.* The estimated concentration change of oxygenated (HbO) and deoxygenated hemoglobin (HbR) at the group level, for each contrast, for session 1 (left) and session 2 (right) separately.

**Table 1.** Group Level Estimated Marginal Means for Contrasts of Interest.

|  | Across Sessions |  | Session 1 |  | Session 2 |  |
| --- | --- | --- | --- | --- | --- | --- |
|  | HbO | HbR | HbO | HbR | HbO | HbR |
| Print Tuning |  |  |  |  |  |  |
| **LIPL** | - | - | .01*<br>[.0031, .016] | - | - | - |
| **LIFG** | - | - | - | - | - | - |
| **RIPL** | .005*<br>[.00048, .0099] | - | .011*<br>[.0038, .0092] | - | - | - |
| Sublexical Tuning |  |  |  |  |  |  |
| **LATL** | - | - | -.009*<br>[-.015, -.0020] | - | .007*<br>[.00011, .013] | - |
| **LIFG** | .01**<br>[.0059, .013] | - | .012**<br>[.0068, .018] | - | .007*<br>[.0018, .012] | - |
| **LIPL** | -.018** | .006* | -.027** | .009* | -.008* | - |
|  | [-.022, -.013] | [.0014, .011] | [-.034, -.021] | [.0020, .015] | [-.015, -.0016] |  |
| LDLPC | - | - | .014*<br>[.0045, .024] |  |  |  |
| RATL | - | - | -.009*<br>[-.015, -.0023] |  |  |  |
| RPTL | - | - | -.008*<br>[-.015, -.0017] |  |  |  |
| RIPL | -.014**<br>[-.019, -.0092] | .007*<br>[.0021, .011] | -.019**<br>[-.025, -.012] | .009*<br>[.0022, .016] | -.009*<br>[-.016, -.0026] | - |
| RIFG | .006*<br>[.0016, .0098] | - | .009*<br>[.0034, .015] | - | - | - |
| Coarse Lexical Tuning |  |  |  |  |  |  |
| **LPTL** | .005*<br>[.00092, .010] | - | - | - | .011**<br>[.0050, .018] | - |
| **LIPL** | .006*<br>[.0016, .011] | - | - | - | .015**<br>[.0080, 0.021] | - |
| LDLPC | .024**<br>[.017, .031] | -.008*<br>[-.014, -.00073] | .022**<br>[.012, .031] | - | .026**<br>[.016, .037] | - |
| RDLPC | -.013*<br>[-.023, -.0040] | - | -.021**<br>[-.034, -.0077] | - | - | - |
| RIFG | -.004*<br>[-.0085, -.00038] | - | - | - | - | - |
| RIPL | - | - | -.013**<br>[-.019, -.0061] | - | - | - |
| RPTL | - | - | -.012**<br>[-.019, -.0055] | - | - | - |
| RATL | - | - | -.007*<br>[-.014, -.00096] | - | - | - |
| LATL | - | - | -.007*<br>[-.014, -.00086] | - | .008*<br>[.0012, .014] | - |
| Fine Lexical Tuning |  |  |  |  |  |  |
| **LIPL** | .024**<br>[.019, .029] | -.008**<br>[-.013, -.0035] | .025**<br>[.019, .032] | -.008*<br>[-.015, -.0015] | .023**<br>[.016, .029] | -.008*<br>[-.015, -.0016] |
| **LATL** | - | - | - | - | - | - |
| LDLPC | .02**<br>[.013, .026] | - | - | - | .032**<br>[.023, .042] | - |
| RDLPC | - | - | -.016*<br>[-.029, -.0027] | - | - | - |
| RIPL | .009**<br>[.0047, .014] | - | - | - | .013**<br>[.0063, .019] | - |
| LPTL | .005* | - | - | - | - | - |
|  | [.00081, .010] |  |  |  |  |  |
| LIFG | -.008** | - | -.012** | - | - | - |
|  | [-.012, -.0046] |  | [-.018, -.0069] |  |  |  |
| RIFG | -.01** | - | -.013** | - | -.007* | - |
|  | [-.014, -.0060] |  | [-.019, -.0071] |  | [-.013, -.0016] |  |

*Note.* This table presents the estimated marginal mean and confidence interval for each contrast of interest, only for the brain regions that were hypothesized to be significantly active for each contrast (listed in bold) and those that were not predicted but identified as significantly active.
\*Significant at $\alpha < 0.05$ , \*\*Significant at $\alpha < 0.001$

Prior to statistical analysis, 8% of runs were removed, due to having more than 8 channels identified as bad (this threshold being >2 SD above the mean number of bad channels across participants). Linear mixed effects analysis was performed on the cleaned data.

### Print Tuning

#### Across Sessions

Results for the contrast consonant strings – false fonts are plotted in Figure 2A. In both the LIPL and LIFG, where a print tuning effect was hypothesized, there was no significant change observed for HbO or HbR. In the RIPL, where a print tuning effect was not hypothesized, there was a significant increase in HbO; this was not associated with a significant decrease in HbR.

#### Sessions 1 and 2

In line with our hypotheses, in the LIPL a significant increase in HbO was obtained in session 1, though this was not associated with a significant decrease in HbR, nor was it replicated in session 2. In the LIFG, where a print tuning effect was hypothesized, there was no significant change observed for HbO or HbR in either session. In the RIPL, consistent with across session results there was a significant increase in HbO observed in session 1, though this was not associated with a significant decrease in HbR, nor was it replicated in session 2.

### Sublexical Tuning

#### Across Sessions

Results for the contrast pseudowords – consonant strings are plotted in Figure 2B. In the LIFG, in line with our hypotheses, there was a significant increase in HbO; a significant decrease in HbR was not obtained. In the LATL, where a sublexical tuning effect was hypothesized, no significant effects were obtained. In the LIPL, where an increase in HbO was predicted, a significant decrease in HbO and significant increase in HbR were obtained. In the RIPL, where a significant effect was not hypothesized, a significant decrease in HbO and significant increase in HbR were obtained. Additionally, in the right inferior frontal gyrus (RIFG), where no effect was predicted, a significant increase in HbO was obtained, though there was no significant decrease in HbR.

#### Sessions 1 and 2

In line with our hypotheses and consistent with across-session results, in the LIFG a significant increase in HbO was obtained in session 1 and 2; this was not associated with a significant change in HbR in either session. In the LATL, where a sublexical tuning effect was hypothesized, there was a significant decrease in HbO observed in session 1, and a significant increase in HbO in session 2; this was not associated with a significant change in HbR in either session. In the LIPL where a significant increase in HbO was hypothesized, a significant decrease in HbO was obtained in both session 1 and 2 (as in the across-session results), with a corresponding significant increase in HbR in session 1 only. In the RIPL, a brain region not hypothesized to demonstrate an effect, a significant decrease in HbO was obtained in session 1 and 2, as well as a significant increase in HbR in session 1 — consistent with the across-session results. Additionally, consistent with across session results, a significant increase in HbO was obtained in the RIFG in session 1, though not in session 2, without a corresponding significant change in HbR. Lastly, in the LDLPC, RATL, and RPTL, brain areas not hypothesized to demonstrate significant sublexical tuning effects, significant changes in HbO were observed in session 1, which were not associated with significant changes in HbR and were not replicated in session 2.

### Coarse Lexical Tuning

#### Across Sessions

Results for the contrast real words – consonant strings are plotted in Figure 2C. In line with our hypotheses, a significant increase in HbO was obtained in the LPTL and in the LIPL; a corresponding decrease in HbR was not significant in either brain region. Additional ROIs not predicted to show effects for this contrast also showed significant effects. In the left dorsolateral prefrontal cortex (LDLPC), a significant increase in HbO and a corresponding significant decrease in HbR were obtained. Additionally, a significant decrease in HbO was obtained in the RDLPC and the RIFG, though these were not associated with significant increases in HbR.

#### Sessions 1 and 2

In the LPTL and LIPL, where a coarse lexical tuning effect was hypothesized and obtained across sessions, a significant increase in HbO was obtained in the session 2 data only; this was not associated with a significant change in HbR for either brain region. In the LDLPC, where an effect was not predicted but observed across sessions, a significant increase in HbO was obtained in session 1 and 2, without a corresponding significant change in HbR in either session. In the RDLPC, where an effect was not predicted but observed across sessions, there was a significant decrease in HbO for session 1, but not session 2, without a corresponding significant change in HbR. In the RIFG, where an effect was not predicted but observed across sessions, no significant changes in HbO or HbR were observed in session 1 or 2. Lastly, while the LATL, RDLPC, RIPL, and RATL did not demonstrate significant effects when data was analyzed across sessions, a significant decrease in HbO was obtained in session 1 for each of these brain regions. Additionally, for the LATL, the reverse effect was observed in session 2, such that a significant increase in HbO was obtained.

### Fine Lexical Tuning

#### Across Sessions

Results for the contrast real words – pseudowords are plotted in Figure 2D. In the LIPL, where an effect was hypothesized, a significant increase in HbO and corresponding significant decrease in HbR was observed. However, in the LATL where an effect was hypothesized, significant changes in HbO or HbR were not obtained. In brain regions not predicted to demonstrate an effect, including the LDLPC, RIPL, and LPTL, significant increases in HbO were obtained. Additionally, in the LIFG and RIFG, where effects were not predicted, a significant decrease in HbO was obtained. These were not associated with significant changes in HbR.

#### Sessions 1 and 2

In the LIPL, in line with our predictions and consistent with across-session results, a significant increase in HbO and corresponding significant decrease in HbR were obtained in session 1 and 2. In the LATL, where a significant effect was also predicted, there were no significant changes in HbO or HbR obtained in session 1 or 2. With respect to ROIs activated in the across-session analysis but not predicted a priori, the LDLPC and RIPL showed a significant increase in HbO in session 2 only. In the LPTL no significant changes in HbO or HbR were obtained in session 1 or 2. In the LIFG, a significant decrease in HbO was obtained in session 1 only. In the RIFG, a significant decrease in HbO was obtained in session 1 and 2. Lastly, in the RDLPC, where a significant effect was not observed in the across-session results, a significant decrease in HbO was obtained in session 1 only.

## Test-Retest Reliability Analysis

To describe test-retest reliability results, we relied on the classification of ICC values defined by Cichetti & Sparrow (1981), such that ICC values are categorized as: excellent (> 0.75), good (0.59 to 0.75), fair (0.40 to 0.58), and poor (< 0.40). Table 2 provides a summary of the significant tuning effects and test-retest reliability results at the ROI level for HbO in select brain areas.

**Table 2.**
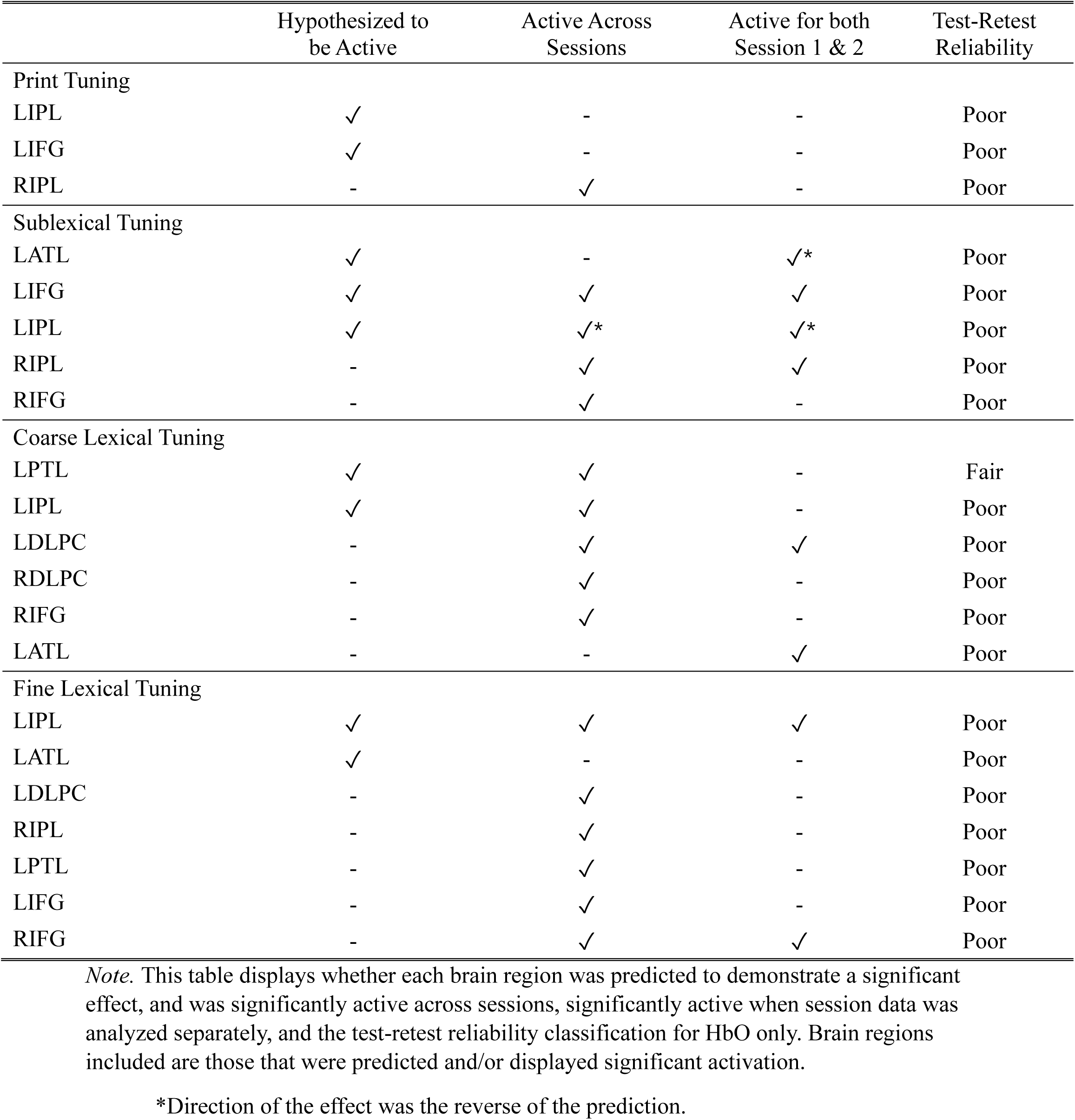
Summary Table of Hypotheses, Active Brain Areas (HbO), and Test-Retest Reliability (HbO)

### Group Level

#### ROI Level

Group level test-retest reliability results for HbO and HbR at the ROI level are presented in Table 3, and are visualized in Supplementary Materials, Figure S5. Overall, we observed poor test-retest reliability in all brain areas hypothesized to be significantly active for print tuning (LIPL and LIFG), sublexical tuning (LATL, LIFG, and LIPL), and fine lexical tuning (LIPL, LATL), despite most of these brain areas being significantly active across sessions (with the exception of the LIFG for print tuning, and the LATL for sublexical and fine lexical tuning) and some being active in sessions 1 and 2. For coarse lexical tuning, we observed fair test-retest reliability in the LPTL for HbO, a region that was hypothesized and significantly active across sessions. However, for the LIPL, also hypothesized and significantly active across sessions for coarse lexical tuning, poor test-retest reliability was obtained.

**Table 3.**
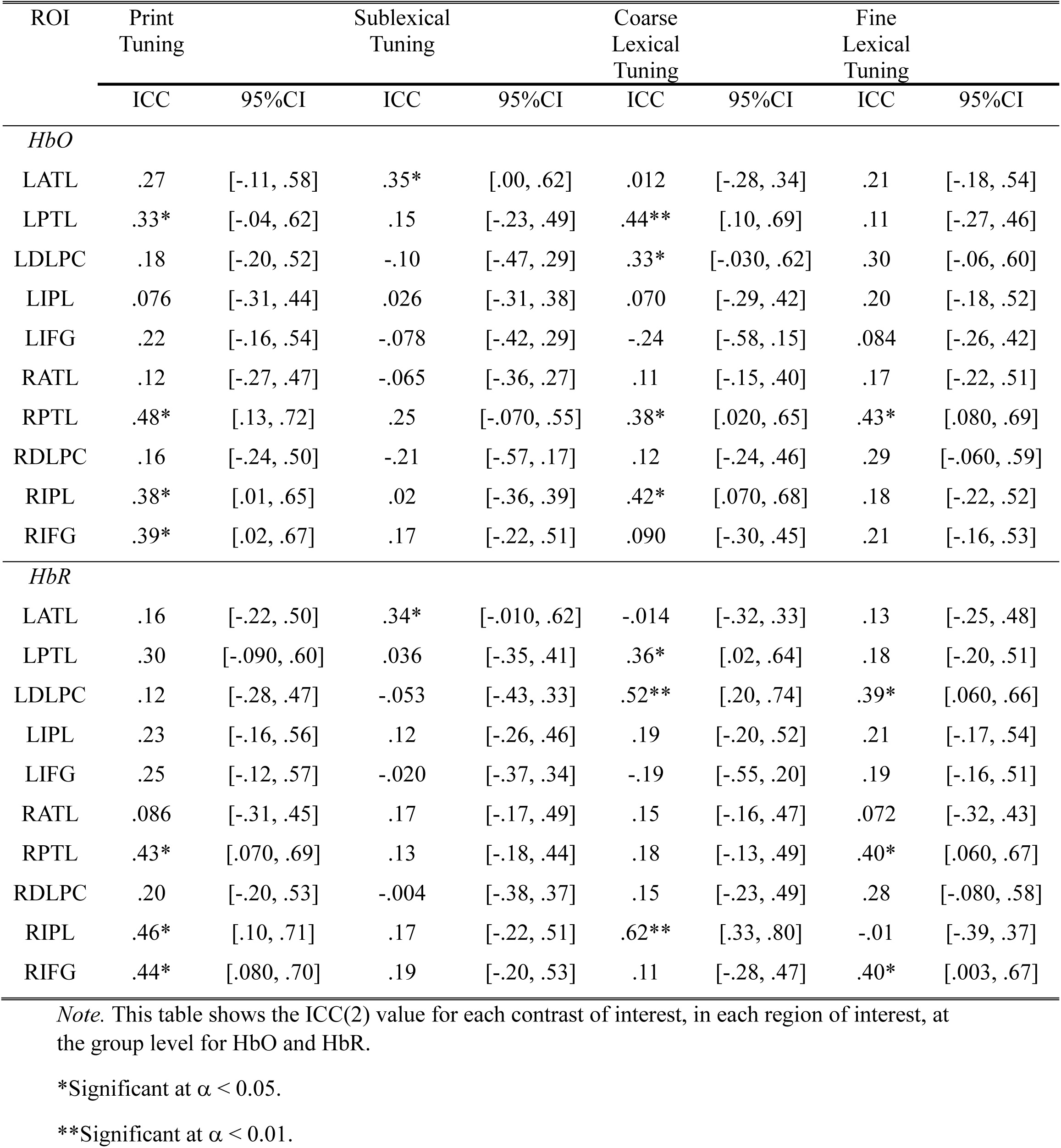
Test-Retest Reliability at the Group Level, by Region of Interest, for HbO and HbR.

| ROI | Print<br>Tuning |  | Sublexical<br>Tuning |  | Coarse<br>Lexical<br>Tuning |  | Fine<br>Lexical<br>Tuning |  |
| --- | --- | --- | --- | --- | --- | --- | --- | --- |
|  | ICC | 95%CI | ICC | 95%CI | ICC | 95%CI | ICC | 95%CI |
| *HbO* |  |  |  |  |  |  |  |  |
| LATL | .27 | [-.11, .58] | .35* | [.00, .62] | .012 | [-.28, .34] | .21 | [-.18, .54] |
| LPTL | .33* | [-.04, .62] | .15 | [-.23, .49] | .44** | [.10, .69] | .11 | [-.27, .46] |
| LDLPC | .18 | [-.20, .52] | -.10 | [-.47, .29] | .33* | [-.030, .62] | .30 | [-.06, .60] |
| LIPL | .076 | [-.31, .44] | .026 | [-.31, .38] | .070 | [-.29, .42] | .20 | [-.18, .52] |
| LIFG | .22 | [-.16, .54] | -.078 | [-.42, .29] | -.24 | [-.58, .15] | .084 | [-.26, .42] |
| RATL | .12 | [-.27, .47] | -.065 | [-.36, .27] | .11 | [-.15, .40] | .17 | [-.22, .51] |
| RPTL | .48* | [.13, .72] | .25 | [-.070, .55] | .38* | [.020, .65] | .43* | [.080, .69] |
| RDLPC | .16 | [-.24, .50] | -.21 | [-.57, .17] | .12 | [-.24, .46] | .29 | [-.060, .59] |
| RIPL | .38* | [.01, .65] | .02 | [-.36, .39] | .42* | [.070, .68] | .18 | [-.22, .52] |
| RIFG | .39* | [.02, .67] | .17 | [-.22, .51] | .090 | [-.30, .45] | .21 | [-.16, .53] |
| *HbR* |  |  |  |  |  |  |  |  |
| LATL | .16 | [-.22, .50] | .34* | [-.010, .62] | -.014 | [-.32, .33] | .13 | [-.25, .48] |
| LPTL | .30 | [-.090, .60] | .036 | [-.35, .41] | .36* | [.02, .64] | .18 | [-.20, .51] |
| LDLPC | .12 | [-.28, .47] | -.053 | [-.43, .33] | .52** | [.20, .74] | .39* | [.060, .66] |
| LIPL | .23 | [-.16, .56] | .12 | [-.26, .46] | .19 | [-.20, .52] | .21 | [-.17, .54] |
| LIFG | .25 | [-.12, .57] | -.020 | [-.37, .34] | -.19 | [-.55, .20] | .19 | [-.16, .51] |
| RATL | .086 | [-.31, .45] | .17 | [-.17, .49] | .15 | [-.16, .47] | .072 | [-.32, .43] |
| RPTL | .43* | [.070, .69] | .13 | [-.18, .44] | .18 | [-.13, .49] | .40* | [.060, .67] |
| RDLPC | .20 | [-.20, .53] | -.004 | [-.38, .37] | .15 | [-.23, .49] | .28 | [-.080, .58] |
| RIPL | .46* | [.10, .71] | .17 | [-.22, .51] | .62** | [.33, .80] | -.01 | [-.39, .37] |
| RIFG | .44* | [.080, .70] | .19 | [-.20, .53] | .11 | [-.28, .47] | .40* | [.003, .67] |

*Note.* This table shows the ICC(2) value for each contrast of interest, in each region of interest, at the group level for HbO and HbR.
\*Significant at $\alpha < 0.05$ .
\*\*Significant at $\alpha < 0.01$ .

Regarding brain regions not hypothesized to be active, for the print tuning contrast, fair test-retest reliability was obtained in the RPTL for HbO, the RIPL for HbR, and the RIFG for HbR. Of these brain areas, only the RIPL was significantly active across sessions, and none were significant at the session level. For the coarse lexical tuning contrast, fair test-retest reliability was obtained in the RIPL for HbO, as well as the LDLPC and RIPL for HbR. Of these brain areas, only the LDLPC was significantly active across sessions, as well as when session data was analyzed individually. For the fine lexical tuning contrast, fair test-retest reliability was obtained in the RPTL for HbO and HbR, and the RIFG for HbR, which were not significantly active across or for individual sessions.

#### Channel Level

At the channel level, we calculated the group level test-retest reliability for HbO and HbR at each channel, for each contrast. These values are shown in Figure 4.

**Figure 4.**
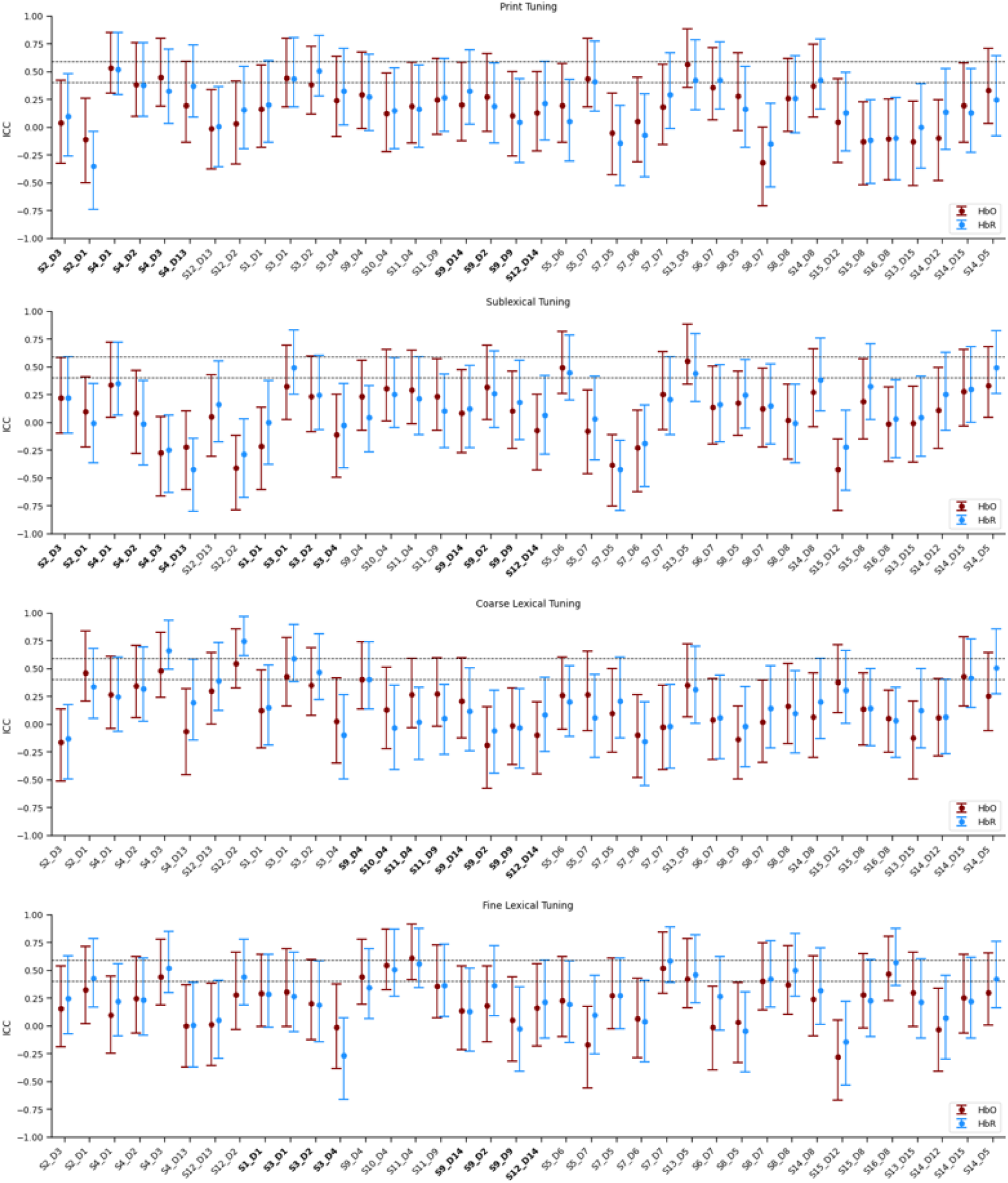
Group-level Test-retest Reliability, at the Channel Level, for HbO and HbR. *Note.* The group level ICC at each channel for both oxygenated (HbO) and deoxygenated (HbR) hemoglobin, for each contrast of interest. Dashed horizontal lines are shown at ICC = 0.40 (i.e., fair test-retest reliability) and ICC = 0.59 (i.e., good test-retest reliability). Channel labels in bold represent channels located within regions of interest that were predicted to demonstrate significant effects for each contrast.

For print tuning, we observed fair test-retest reliability in 5 of 38 channels measured for HbO, and in 7 of 38 channels for HbR, with poor test-retest reliability in the remaining channels. Of the channels demonstrating fair reliability for HbO and HbR, one channel covered the LIFG, a hypothesized brain region that did not demonstrate a significant print tuning effect. This channel demonstrated fair reliability for HbO (ICC = .41, CI95% = [.04, .67], *F* = 2.35, *p* = .015) and HbR (ICC = .41, CI95% = [.04, .68], *F* = 2.34, *p* = .016). All other channels that obtained fair test-retest reliability were included in ROIs not predicted to demonstrate significant print tuning effects.

For sublexical tuning, 2 of 38 channels for HbO and 4 of 38 channels for HbR obtained fair test-retest reliability, with poor test-retest reliability in the remaining channels. Of the channels demonstrating fair test-retest reliability for HbO and HbR, one channel was measuring the LATL, a hypothesized ROI that did not demonstrate a significant sublexical tuning effect. This channel demonstrated fair reliability for HbO (ICC = .50, CI95% = [.18, .73], *F* = 3.31, *p* = 0.001) and HbR (ICC = .45, CI95% = [.12, .70], *F* = 2.83, *p* = .004). Additionally, one channel measuring the LIPL demonstrated fair test-retest reliability for HbR (ICC = .45, CI95% = [.11, .70], *F* = 2.68, *p* = .006). The LIPL was predicted and was significantly active both across sessions and in session 1 and 2 separately. All other channels that obtained fair test-retest reliability were included in ROIs not predicted to demonstrate significant sublexical tuning effects.

For coarse lexical tuning, 6 of 38 channels for both HbO and HbR demonstrated fair or higher test-retest reliability. For HbO, two channels measuring the LPTL demonstrated fair test-retest reliability (ICC = .47, CI95% = [.08, .72], *F* = 3.47, *p* < .001; ICC = .43, CI95% = [.070, .69], *F* = 2.49, *p* = .011). The LPTL was predicted and demonstrated a significant coarse lexical tuning effect across sessions for HbO. For HbR, of the channels demonstrating fair test-retest reliability, one channel measured the LPTL (ICC = .42, CI95% = [.070, .68], *F* = 2.46, *p* = .011) and one channel the LIPL (ICC = .47, CI95% = [.12, .72], *F* = 2.73, *p* = .0057), which were both predicted brain regions but only demonstrated significant coarse lexical tuning effects for HbO. All other channels that obtained fair or higher test-retest reliability were in ROIs not predicted to demonstrate significant coarse lexical tuning effects.

For fine lexical tuning, 8 of 38 channels for HbO and 10 of 38 channels for HbR demonstrated fair test-retest reliability. For HbO and HbR, none of the channels demonstrating fair test-retest reliability were measuring from brain regions predicted to demonstrate a significant fine lexical tuning effect.

### Subject Level

Two subjects’ data were not analyzed at the subject level, as the effect size values were too close to zero for the ICC analysis; thus, the following analysis includes 28 subjects. Test-retest reliability within individual participants was highly variable at the ROI level for both HbO and HbR. Notably, for each contrast, fewer than 50% of subjects demonstrated test-retest reliability that was fair or higher in predicted ROIs. Subject level test-retest reliability for HbO and HbR within each ROI is presented using box plots in Figure 5, to provide a visualization of the range across subjects.

**Figure 5.**
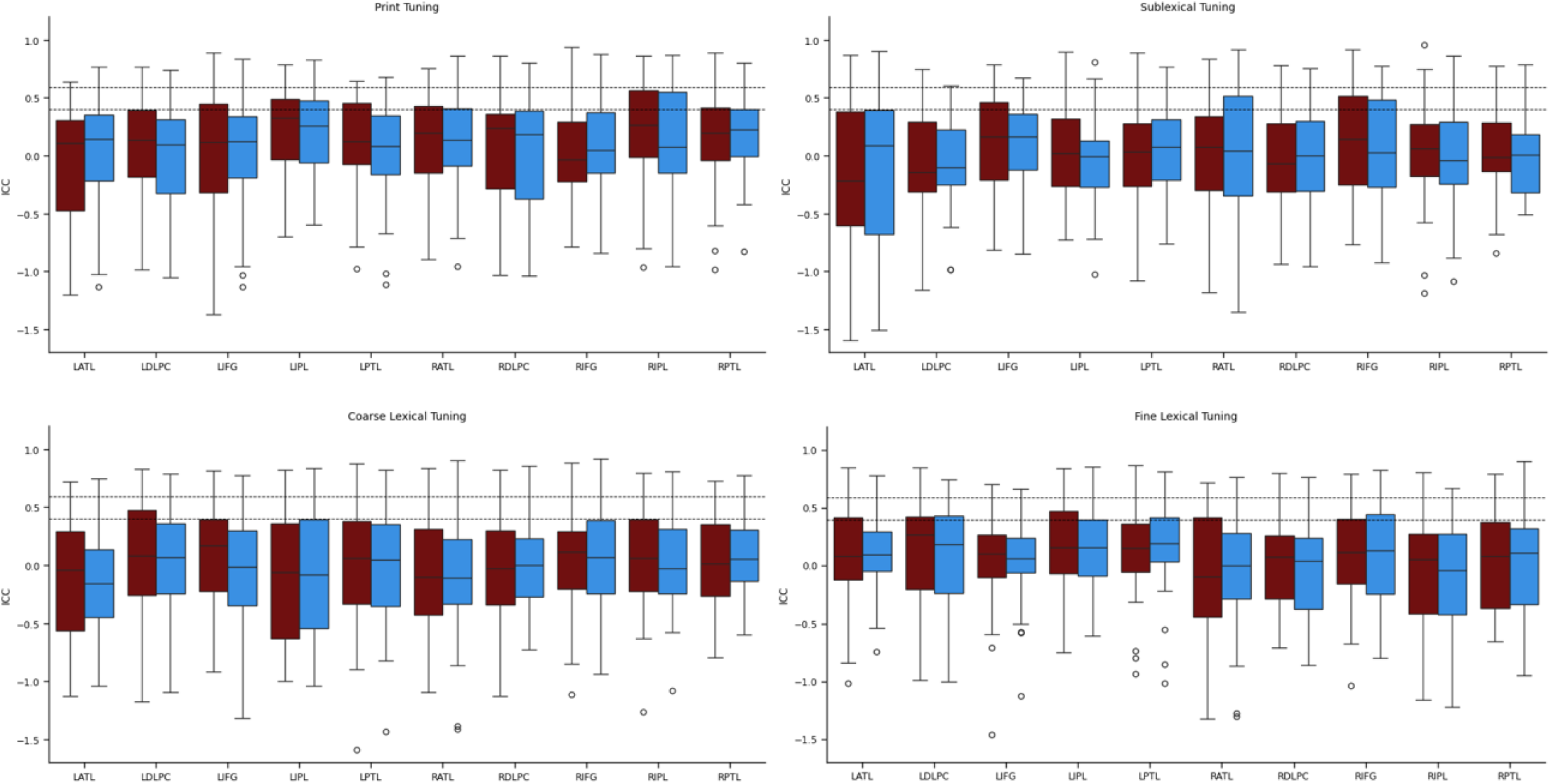
Subject Level Test-Retest Reliability, at the Region of Interest Level, for HbO and HbR. *Note.* The mean ICC value across individual subjects at each brain region, plotted for oxygenated (HbO; maroon) and deoxygenated (HbR; blue) hemoglobin for each contrast of interest. Dashed horizontal lines are shown at ICC = 0.40 (i.e., fair test-retest reliability) and ICC = 0.59 (i.e., good test-retest reliability).

#### Print Tuning

Across ROIs for the print tuning contrast, subjects demonstrated test-retest reliability ranging from poor to excellent for HbO and HbR. When considering predicted ROIs, average test-retest reliability in the LIPL across subjects was poor for HbO (*M* = .21, *SD* = .37) and HbR (*M* = .24, *SD* = .32), but ranged from poor to excellent across subjects for HbO and HbR. Similarly, in the LIFG, average test-retest reliability across subjects was poor for HbO (*M* = .13, *SD* = .47) and HbR (*M* = -.011, *SD* = .52), and ranged from poor to excellent across subjects for HbO and HbR.

#### Sublexical Tuning

Across ROIs for the sublexical tuning contrast, subjects demonstrated test-retest reliability ranging from poor to excellent for HbO and HbR. When considering only predicted ROIs, average test-retest reliability across subjects in the LATL was poor for HbO (*M* = -.11, *SD* = .63) and HbR (*M* = -.13, *SD* = .65), and ranged from poor to excellent for HbO and HbR. In the LIFG, average test-retest reliability across subjects was also poor for HbO (*M* = .088, *SD* = .50) and HbR (*M* = .043, *SD* = .43), and ranged from poor to excellent for HbO and poor to good for HbR. Lastly, in the LIPL, average test-retest reliability across subjects was poor for HbO (*M* = .026, *SD* = .44) and HbR (*M* = -.058, *SD* = .44), and ranged from poor to good for HbO and HbR.

#### Coarse Lexical Tuning

Across ROIs for the coarse lexical tuning contrast, subjects demonstrated test-retest reliability ranging from poor to excellent for HbO and HbR. Considering only predicted ROIs, average test-retest reliability across subjects in the LPTL was poor for HbO (*M* = .018, *SD* = .49) and HbR (*M* = .032, *SD* = .37), and ranged from poor to excellent for HbO and HbR. In the LIPL, average test-retest reliability across subjects was poor for HbO (*M* = - .055, *SD* = .54) and HbR (*M* = -.0010, *SD* = .52), and ranged from poor to excellent for HbO and HbR.

#### Fine Lexical Tuning

Across ROIs for the fine lexical tuning contrast, subjects demonstrated test-retest reliability ranging from poor to excellent for HbO and HbR. Considering only predicted ROIs, average test-retest reliability across subjects in the LIPL was poor for HbO (*M* = .18, *SD* = .37) and HbR (*M* = .17, *SD* = .36), and ranged from poor to good for HbO and poor to excellent for HbR. In the LATL, average test-retest reliability across subjects was poor for HbO (*M* = .063, *SD* = .46) and HbR (*M* = .081, *SD* = .36), and ranged from poor to excellent for HbO and HbR.

### Subject Level Test-Retest Reliability and Behavioral Data

Linear mixed effects analysis was performed to explore the effect of the difference of each relevant behavioral variable from session 1 to session 2 on test-retest reliability at the subject-level. Regarding the PANAS-X indices (negative affect, positive affect, fatigue, and attentiveness index), there were no significant main effects or interactions predicting subject level test-retest reliability. Similarly, for sleep quantity, sleep quality, and caffeine consumption, there were no significant main effects or interactions predicting subject level test-retest reliability.

It is important to note these analyses were exploratory, as this study was not adequately powered to detect the presence of statistically significant main effects or interactions on the test-retest reliability data. However, the relatively small F-values observed provide little support to suggest these variables meaningfully influence test-retest reliability at the subject level.

## Discussion

The present study assessed the test-retest reliability of the fNIRS signal during a lexical decision task in the left and right hemispheres of the inferior frontal, superior temporal, and inferior parietal cortices. We analyzed the location, magnitude, and reliability of the fNIRS signal for common lexical decision effects. In general, when we collapsed data across sessions, each contrast of interest yielded significant activation (as indexed by HbO) in at least one predicted ROI, and activation in these ROIs was largely replicated when session date was analyzed separately. At the same time, however, at least one predicted ROI for each contrast did not show significant activation across or within sessions. As well, test-retest reliability across most regions and contrasts was poor at the group level, although it ranged from poor to excellent within individual participants. Most of our behavioral predictors, including affect, sleep, and caffeine consumption did not differ significantly between sessions, nor did any show a statistical relationship with ICC values, suggesting these variables were not responsible for the generally poor test-retest reliability of the measurements. Below we consider these findings in more detail.

### Behavioral Data

We did not observe a significant difference in participants’ negative affect or fatigue, as assessed by the PANAS-X,^56^ or their sleep quantity, sleep quality, or caffeine consumption prior to each study session. There was a significant decrease in positive affect and attentiveness observed at session 2, as assessed by the PANAS-X.^56^ However, changes in positive affect have been primarily associated with changes in brain activity in medial frontal and subcortical brain areas of the limbic system,^70,71^ suggesting this would have minimal effect on task-related activation in ROIs in the present study. Additionally, while changes in attentiveness may influence participants’ task performance, we did not observe a decrease in accuracy or reaction time at session 2.

Regarding behavioral performance on the lexical decision task, all participants demonstrated near-perfect accuracy and fast reaction times for each word type in both sessions 1 and 2. Further, there was no significant difference between group level accuracy for session 1 and 2 for most word types, though accuracy for real words improved in session 2, and reaction times were faster for all word types at session 2. This suggests participants demonstrated similar effort on the task across sessions, though practice effects may have improved their accuracy and response speed in session 2. Practice effects typically result in a change in performance across all task conditions equally,^72^ which is in line with the change in reaction time across sessions. The observed increase in accuracy for real words may have been due to the increased number of presentations of each real word, as each real word was presented three times in each session, and the same words were used in both sessions. Therefore, by session 2, participants had the most exposure to the real words, which may have led to more accurate recognition.

### fNIRS Data

fNIRS data was first analyzed across sessions, to obtain the most sensitivity for identifying significant effects. Considering first the print tuning contrast, we hypothesized increased activation would be observed in the LIPL and LIFG for consonant string as compared to false font stimuli. In line with our hypotheses, we observed significant activation in the LIPL (increased HbO), however this significant effect was only maintained in session 1 when session data was analyzed separately. Additionally, and contrary to our hypotheses, we did not observe a significant print tuning effect in the LIFG. Broadly, print tuning is defined as the increased neural sensitivity to text stimuli, and is well documented in the fMRI literature as occurring in the LIPL and LIFG, as well as regions not covered by our fNIRS montage.^34–36^ Therefore, while it is promising to observe significant activation in the LIPL, the lack of significant activation in the LIFG is not in line with prior fMRI research.

For the sublexical tuning contrast, defined as the difference between pronounceable non-words and consonant strings, we observed the predicted increase in HbO in the LIFG across-sessions, and this effect was significant when session data was analyzed individually. However, we did not observe the predicted effects in the LIPL or LATL. This is only partially consistent with the fMRI literature, where sublexical tuning (and related contrasts involving converting orthographic to phonologic information) has been consistently associated with increased activation not only in the LIFG,^39–41^ but also the LIPL^39^ and LATL^37^. It is notable, however, that the fNIRS data distinguish the print tuning from the sublexical tuning conditions by eliciting significant activation in different brain regions.

For coarse lexical tuning, defined as the difference between real words and consonant strings, we observed results that supported our hypotheses. Specifically, we observed a significant increase in HbO in both the LPTL and LIPL, consistent with the fMRI literature,^42,43^ though these when data was analyzed at the individual session level this was only sustained in session 2 data.

Lastly, for fine lexical tuning, our hypotheses were partially supported. In the LIPL we observed a significant increase in HbO and significant decrease in HbR, present across sessions and in each session individually. Increased neural activity in the LIPL for fine lexical tuning has been documented in both fMRI literature^44–49^ and in a prior lexical decision study employing fNIRS.^33^ However, we did not observe the predicted effect in the LATL, thus failing to replicate prior fMRI studies that have reported lexical tuning effects.^44,45,49^

Overall, across contrasts of interest, we observed multiple significant contrast-related changes in HbO in brain regions consistent with prior fMRI literature, when data was averaged across two sessions of data collection. However, these significant results were not maintained for all brain regions when data was analyzed at the single session level. Moreover, other areas predicted from the fMRI literature were not identified as active across sessions, or at the individual session level. This does not appear simply to be a lack of fNIRS sensitivity to specific brain areas; for example, both LIFG and LIPL activation were predicted for the print and sublexical tuning contrasts, but we obtained only LIPL for print tuning and only LIFG for sublexical tuning. Instead, these results may point to a limited sensitivity of fNIRS more broadly, particularly when investigating contrasts that result in a relatively small magnitude change in activation (e.g., comparing pseudowords to strings of consonants). In line with this interpretation, across brain regions, the significant contrast-related increases in HbO were often not associated with a corresponding significant decrease in HbR. Changes in HbO are typically higher in amplitude and have a higher signal-to-noise ratio than HbR, making HbO more robust when the change in activation is smaller or signal strength is weaker.^2^ Thus, while fNIRS is adequately sensitive to detect reading-related brain activity during lexical decision, its sensitivity may be limited by the magnitude of the contrast being investigated.

### Test-Retest Reliability

We hypothesized we would observe test-retest reliability in the range of good (0.59 < ICC < 0.75) at the group level. However, we observed consistently poor test-retest reliability when analyzing group data in the hypothesized ROIs for each contrast and in brain areas that demonstrated significant task-related activation. The low ICC values are particularly striking for brain regions that demonstrated significant changes in HbO and HbR both across sessions and in session 1 and 2 separately. That is, the activation was consistent from one session to the next, but the ICC results for those regions were still poor.

When analyzing group level test-retest reliability at the channel level, we identified some channels that demonstrated fair to excellent test-retest reliability. However, the majority of these sampled from brain regions not predicted (or demonstrated) to show significant effects; this may simply reflect channels that reliably showed a lack of task-related changes.

Additionally, we had hypothesized we would observe fair test-retest reliability across participants at the subject level, but on average subjects demonstrated poor test-retest reliability in brain regions predicted to demonstrate significant effects. While test-retest reliability of the fNIRS signal ranged from poor to excellent across subjects in predicted brain regions, most subjects (>50%) demonstrated poor test-retest reliability across contrasts, consistent with the average. These results are somewhat in line with existing literature, as prior investigations of the test-retest reliability of the fNIRS signal have documented high variability in subject level reliability, and task-related activation at the subject level.^4,9,18^

Overall, test-retest reliability at the group and subject level was lower than predicted, and group level results were lower than what has been reported in prior literature. While existing studies of the test-retest reliability of fNIRS at the group level have similarly reported varying test-retest reliability across ROIs,^9,17–19^ most have still obtained fair,^19^ or good to excellent^4,9,15,16,18^ test-retest reliability in predicted ROIs. The most notable difference between the present study and existing literature is prior work has utilized slower event-related (e.g., Plichta et al., 2006), or block design (e.g., de Rond et al., 2023; Dong et al., 2024; Huang et al., 2017; Ranchet et al., 2023; Schecklmann et al., 2008; Wiggins et al., 2016) paradigms. In the present study, words were only presented for 2 s, separated by short inter-trial intervals (5 s), which does not provide adequate time for the hemodynamic response to fully return to baseline prior to the next stimulus (often from a different condition). This ultimately results in a signal that is smaller and harder to deconvolve than if the hemodynamic response was able to fully return to baseline between trials (i.e., slow event-related design), or summated across multiple stimuli of the same type (i.e., block design).^3^ Additionally, this study focused on relatively small differences between stimulus conditions (e.g., consonant strings – pseudowords), which similarly reduces the magnitude of the change in hemodynamic response. Therefore, it is possible fNIRS may be more likely to demonstrate adequate test-retest reliability for tasks designed to maximize the measured hemodynamic response (e.g., block design) and that investigate more robust contrasts (e.g., stimulation compared to baseline).

In addition, it is still unclear what variables are driving subject level differences in reliability. While we measured certain variables that might affect reliability — including caffeine consumption, sleep quantity, sleep quality, participant fatigue, and affect — none of these showed a significant relationship with ICC values. Another variable that has previously been hypothesized to affect fNIRS reliability is optode placement.^19,73^ Specifically, from one session to the next, the placement of the fNIRS cap holding the optodes may vary. While we employed the well-established International 10-10 system for fitting the cap on the head, this has limitations. Notably, the stretchy nature of the cap, combined with how hair may be distributed under the cap, can be expected to produce some variation in the position of optodes each time the cap is placed on the head. In line with this, de Rond et al. (2023) observed significantly lower fNIRS test-retest reliability when participants removed the cap between measurement instances than when repeated measurements were obtained without removing the cap. Notably, in the present study, we did not have equipment capable of quantifying the variance in cap placement and so it is possible that, at the individual level, the set of channels covering an ROI may have covered active cortex to a greater or lesser extent from one session to the next.

Another consideration is that while fMRI studies perform spatial normalization to align the size and complex cortical folding patterns of individuals into a common space, this is not routinely done in fNIRS. Doing so would require technology to measure optode placement at each session with high precision (e.g., photogrammetry) equipment that most fNIRS labs lack— as well as anatomical MRI scans of each individual’s brain, which somewhat undermines the advantages of using fNIRS over fMRI. Therefore, it is possible a specific fNIRS montage accurately samples from active brain regions in some participants, but not others, simply due to variance in their underlying neuroanatomy and idiosyncratic brain organization.

## Future Directions

The present study aimed to identify the test-retest reliability of the fNIRS signal during a lexical decision task, to support future use of fNIRS to investigate individual differences in brain activation and how that relates to behavioral measures of reading skill. Considering the high level of variability in both task-related activation and test-retest reliability identified at the individual level, the results of this study suggest fNIRS may not be suitable for investigating individual differences in brain activation during a fast event-related lexical decision task. Further work is needed to determine how best to overcome the factors contributing to its low reliability. One avenue for investigation is the effects of task design; for example, block design studies may yield more reliable activation by virtue of having higher contrast-to-noise ratios. Another is using techniques such as photogrammetry to quantify variance in optode location across sessions and provide guidance to minimize this variance, as well as perhaps integrating any residual variance into the analysis pipeline (e.g., through some form of spatial normalization).

## Conclusions

Overall, the present findings provide evidence suggesting fNIRS is adequately sensitive to detect reading-related brain activity during lexical decision at the group level. Moreover, the present study demonstrated that the ROIs showing significant task-related brain activity measured by fNIRS largely replicated across sessions at the group level, despite poor test-retest reliability values. Notably, it is relevant to consider the sensitivity of fNIRS to task-related brain activity may be limited by the magnitude of the contrast being investigated, as well as the coverage and consistency of coverage of specific brain regions across participants. Ultimately, to continue advancing the use of fNIRS in the study of reading, the field will need to develop protocols for ensuring precision and consistency of optode placement both across participants and measurement instances.

## Data and Code Availability

The data presented in this article, as well as the analysis code, are publicly available on Borealis in the repository “Investigating fNIRS Test-Retest Reliability During Lexical Decision” at https://doi.org/10.5683/SP4/MXMU9I.

## Acknowledgements

The authors would like to acknowledge the contributions of Dr. Carrie Demmans Epp and Gisele Arevalo for their development of the pseudowords used in the lexical decision task. This research was supported by an Insight Grant (435-2020-1471) from the Social Sciences and Humanities Research Council (SSHRC) of Canada, awarded to AJN, as well as a SSHRC Partnership Grant (895-2020-1004; J.F. Werker, PI). LE was supported by a SSHRC Canada Graduate Scholarship (Master’s Level), Nova Scotia Graduate Scholarship (Doctoral), and Dalhousie Centre for Psychological Health Bursary. SR was supported by a Killam Predoctoral Scholarship Level 2 (Doctoral), SSHRC Doctoral Fellowship, and Nova Scotia Graduate Scholarship (Doctoral).

