## Supplemental figures and tables for "Investigating fNIRS Test-Retest Reliability During Lexical Decision"

### 1. SUPPLEMENTARY RESULTS

#### 1.1 Behavioural Analyses

**Figure S1**

*Group-level PANAS-X Index Scores by Session*

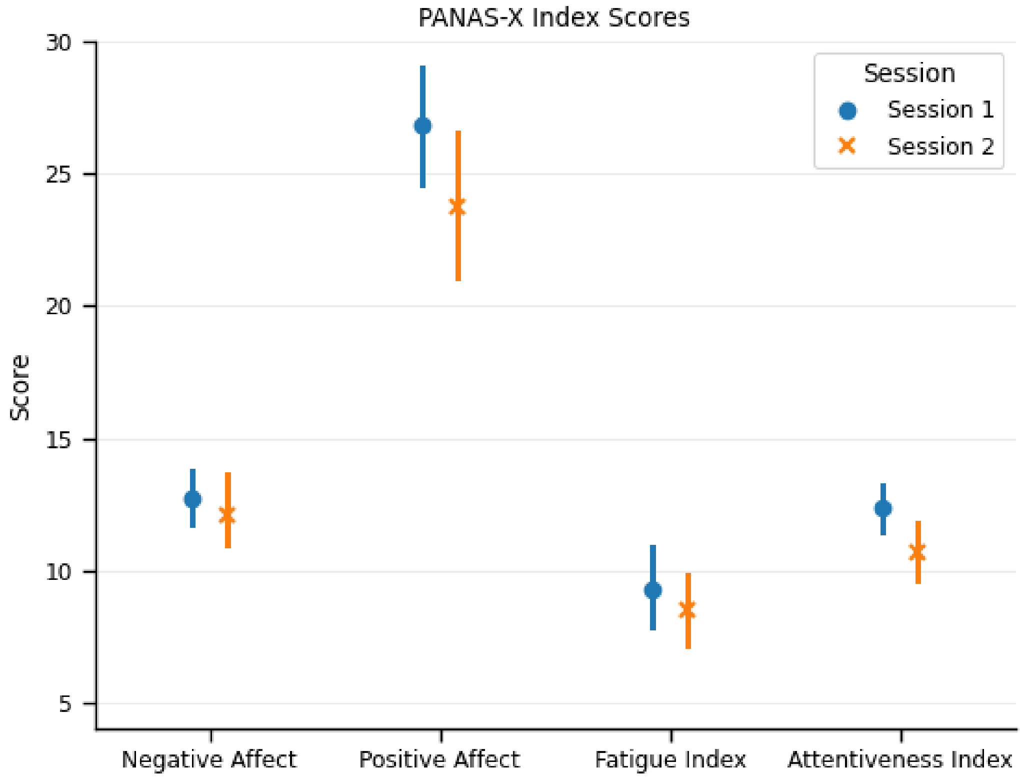


*Note.* Visualization of the group mean scores, and 95% confidence intervals around the mean, on the relevant indices of the PANAS-X (Watson & Clark, 1994) for session one and two.

**Figure S2**

*Self-Reported Sleep Quantity, Sleep Quality and Caffeine Consumption by Session*

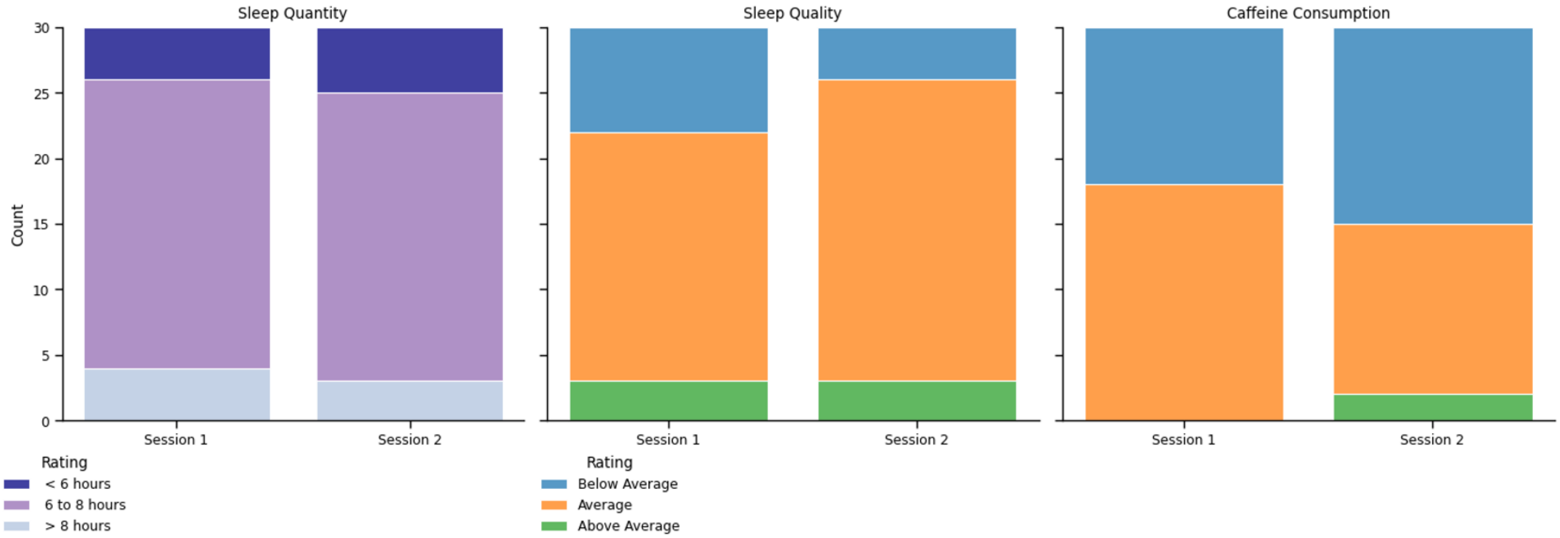


*Note.* Visualization of self-reported sleep quantity, sleep quality and caffeine consumption, for session one and two. Participants rated sleep quality and caffeine consumption based what is typically average for themselves, for each variable.

**Figure S3**

*Group Level Accuracy on the Lexical Decision Task*

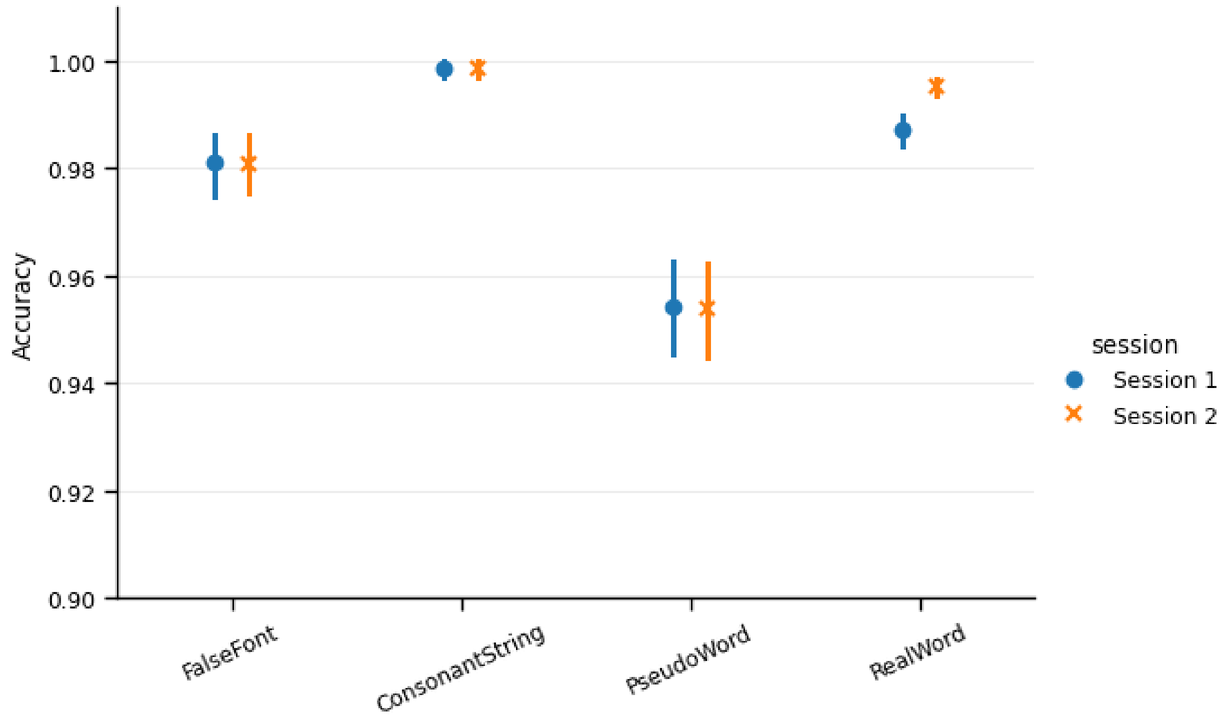


*Note.* Visualization of the group mean accuracy, and 95% confidence intervals around the mean, for each word type presented during the lexical decision task, for session one and two.

**Figure S4**

*Group Level Reaction Time on the Lexical Decision Task*

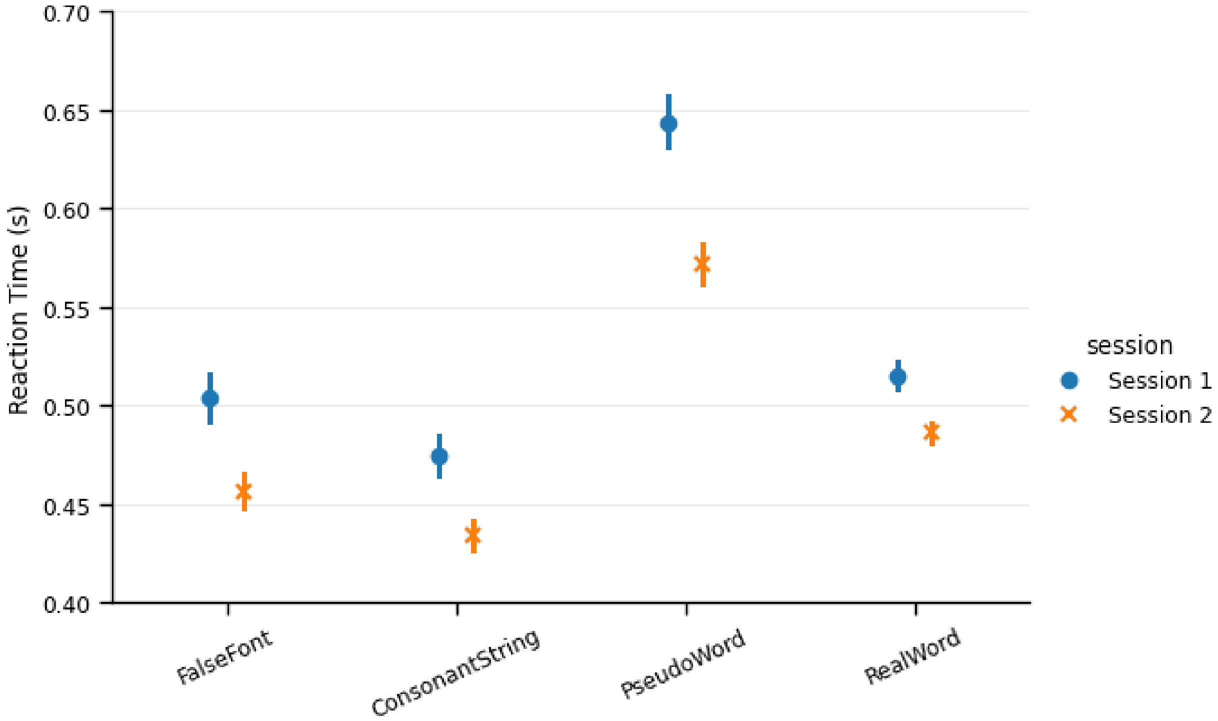


*Note.* Visualization of the group mean reaction time, and 95% confidence intervals around the mean, for each word type presented during the lexical decision task, for session one and two.

#### 1.2 Test-Retest Reliability Analyses

**Figure S5**

*Group Level Test-Retest Reliability, at the Region of Interest Level, for HbO and HbR*

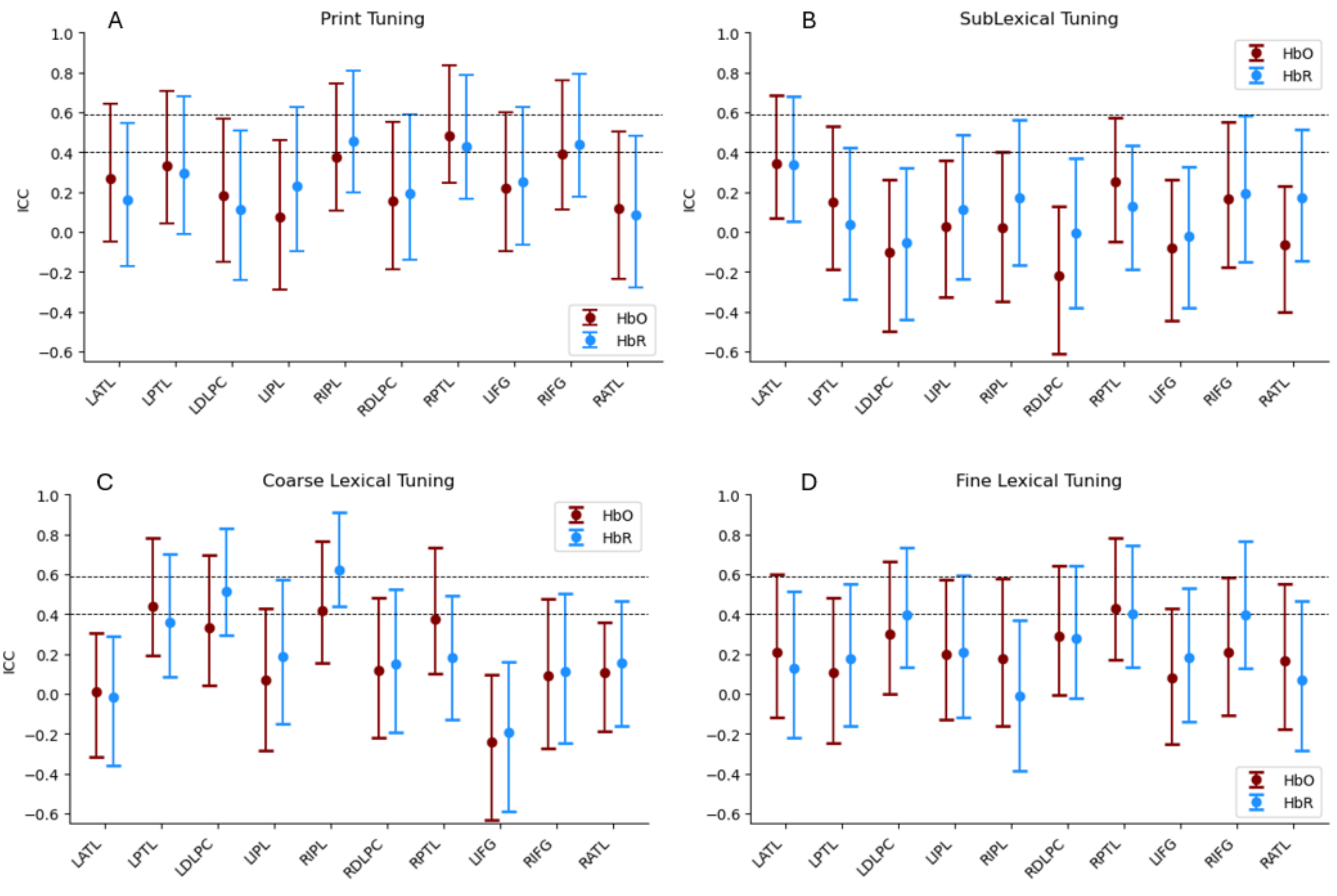


*Note.* The ICC values analyzed across subjects at the region of interest level for both oxygenated (HbO) and deoxygenated (HbR) hemoglobin, for (A) print tuning, (B) sublexical tuning, (C) coarse lexical tuning, and (D) fine lexical tuning contrasts. Dashed horizontal lines are shown at ICC = 0.40 (i.e., fair test-retest reliability), and ICC = 0.59 (i.e., good test-retest reliability).
